# hMSCs form Osteocyte-like Cell Networks within Strain-Stiffening Bottlebrush Polymer Hydrogels

**DOI:** 10.64898/2026.09.22.753603

**Authors:** Monica L. Ohnsorg, Nicole E. Friend, Sophie E. Givens, Daniel Saeb, Kayla M. Mash, Kristi S. Anseth

## Abstract

Bone formation and remodeling depend on dynamic biochemical and biomechanical signaling from the collagen-rich osteoid that precedes mineralization, yet the role of osteoid nonlinear mechanics in regulating osteocyte-like network formation remains poorly understood. Here, we engineered a synthetic bottlebrush polymer hydrogel (BB) that mimics key mechanical features of osteoid and compared it to collagen type-I (Col1) matrices with matched shear modulus (∼70 Pa) and strain-stiffening behavior. Human bone marrow-derived mesenchymal stem/stromal cells (hMSCs) were cultured for 28 days in growth (GM) or osteogenic media (OM) to examine network formation, and functional connectivity using live cell fluorescence recovery after photobleaching. hMSCs cultured in BB networks and OM showed upregulation of early osteocyte markers compared to Col1. We find that strain-stiffening materials with minimal stress relaxation promote osteocyte-like cell differentiation with functional connectivity, establishing osteoid-mimetic BB hydrogels as a promising matrix to study human osteocytogenesis and osteocyte mechanotransduction *in vitro*.

## Introduction

Bone formation and remodeling is a dynamic, multistep process that depends not only on biochemical signaling but also on the evolving material properties of the extracellular matrix (ECM). During intramembranous ossification, human bone marrow–derived mesenchymal stromal cells (hMSCs) differentiate into osteoprogenitor cells and osteoblasts that secrete a provisional matrix known as bone osteoid ^1^. The osteoid is compositionally distinct from mature bone, consisting of approximately 90% type-I collagen and 10% non-collagenous proteins^2,3^. This microenvironment supports osteoblast embedding and a subsequent phenotypic transition resulting in cellular network formation and differentiation into osteoid osteocytes prior to mineralization. The collagen fibrils support the deposition and organization of calcium phosphate crystals, in the form of hydroxyapatite around the network of connected osteocytes^3–5^. During bone remodeling, bone resorption by osteoclasts is followed by deposition of a collagen-rich osteoid by mesenchymal cells and osteoblasts. As the cells are embedded within the osteoid, they form connected networks through dendritic protrusions indicative of pre-osteocytic and mature osteocyte phenotypes before mineralized bone is re-established^6^. Despite the central role of this collagen-rich secreted ECM in regulating bone quality during both bone formation and regeneration^5,7,8^, the contribution of biophysical matrix signals, especially non-linear biomechanics, and stimuli that guide these dramatic changes in osteogenic cell phenotype remain poorly understood. Thus, synthetic extracellular materials provide an opportunity to study how osteoid mechanics regulate cell morphology and connectivity *in vitro* to begin to develop methods for hypothesis testing that can complement *in vivo* models of bone development, injury, and disease.

To date, models of intramembranous ossification *in vitro* have been inherently challenging due to the simultaneous and continuous changes in ECM composition, structure, and mechanics that occur over the differentiation timeline. Traditional two-dimensional culture systems fail to capture the three-dimensional context required for osteocyte differentiation, while many existing three-dimensional models prioritize mineral deposition rather than the earlier stages of cell embedding and network formation ^9^. As a result, there is an unmet need for 3D *in vitro* models of bone development and bone remodeling that can support osteogenic differentiation while preserving the hallmark dendrite-rich morphology and functional connectivity characteristic of osteocyte-like cells.

Several innovative approaches have partially addressed this challenge. Microporous, degradable poly(ethylene glycol) (PEG) scaffolds have been synthesized via polymerization induced phase separation of dextrose and hyaluronic acid to form a void-rich PEG hydrogel. These scaffolds supported osteogenic differentiation of hMSCs and the expression of early osteocyte markers, such as E11, along with protrusion-rich cell morphologies^10^. In an alternative approach, interpenetrating networks composed of alginate and collagen with tunable stress-relaxation characteristics enabled three-dimensional culture of a murine osteocyte-like cell line, IDW-SD3, and results demonstrated some limited dendritic connections and functional gap junction formation between adjacent cells^11^. Finally, IDW-SD3s embedded in MMP-degradable PEG hydrogels were found to undergo an osteoblast-like to osteocyte-like transition more readily than when cultured in nondegradable hydrogels or on 2D surfaces^12^. Similarly, OCY454 osteocyte-like cells cultured within 3D porous scaffolds increased expression of bone remodeling factors compared to 2D culture^13^. Several other natural scaffolds like modified collagen and silk fibroin have also been used to support the osteogenic differentiation of hMCSs^14,15^. Collectively, these contributions underscore the importance of 3D context, but thus far, the ability to robustly and reproducibly support the formation of a connected and percolating osteocyte-like cellular network, as observed *in vivo* has been limited.

One key feature of the osteoid ECM that has been underrepresented in many synthetic culture platforms is its distinct mechanical behavior. Collagen-rich matrices exhibit nonlinear elastic properties, including strain-stiffening, as well as time-dependent stress-relaxation behavior^16,17^. The interplay between these two time-dependent biomechanical responses arise from the semi-flexible fibrillar architecture of collagen-rich ECM with strain-stiffening or nonlinear elasticity timescales that relate to local fiber alignment on the order of minutes and stress-relaxation or viscoelastic behavior occurring due to ECM rearrangement over the course of hours^18^. As a result, both are often important regulators of cell mechanotransduction. For example, synthetic fibrillar and crosslinked hydrogels, which recapitulate strain-stiffening mechanics similar to collagen and fibrin, have been recently highlighted as promising tools for studying cell–matrix interactions in three dimensions^16^. In particular, recent work demonstrated that synthetic bottlebrush polymer hydrogels can be engineered to exhibit strain-stiffening behavior within the biologically relevant stiffening regime (BRSR, < 25 Pa), leading to pronounced protrusion-rich morphologies in hMSCs driven by direct cell– matrix interaction^19^. However, the relevance of such nonlinear elastic mechanics to osteogenic differentiation and osteocyte network formation, particularly in relation to the collagen-rich osteoid, has not been systematically examined.

Motivated by these prior findings, we sought to engineer a three-dimensional *in vitro* platform that captures key mechanical features of collagen-rich osteoid while enabling systematic interrogation of osteogenic cell behavior during early stages of intramembranous-like ossification. We optimized a synthetic bottlebrush polymer hydrogel (BB) to match the modulus and strain-stiffening response of type I collagen hydrogels (Col1), while intentionally minimizing stress-relaxation behavior. By directly comparing bottlebrush polymer hydrogels and collagen hydrogels with similar initial stiffness and nonlinear elasticity but distinct viscoelastic profiles, we aimed to isolate the role of strain-stiffening biomechanical cues in regulating osteogenic differentiation and network formation.

Furthermore, while bottlebrush polymer hydrogels have previously been shown to induce spindle-like morphologies in short-term hMSC culture^19^, we significantly extended the culture duration to four weeks, introduced defined biochemical stimuli, and examined how nonlinear elastic ECM cues influenced cell behavior over time. Through this approach, we address a central biological challenge in *in vitro* models of bone regeneration: the lack of a three-dimensional material microenvironment that reliably supports the formation and function of connected osteocyte-like cell networks. Results herein focus on examining the material properties of a collagen-rich osteoid, rather than the terminal mineralized matrix, and reveal insight as to how ECM mechanics can regulate critical transitions in early bone regeneration and remodeling. Fluorescence recovery after photobleaching (FRAP) measurements on live cells in 3D were used to validate cell network connectivity. The role of gap junctions in these cell network connections was investigated through immunocytochemical staining and quantitative image analysis of connexin 43 followed by further FRAP measurements after dosing with a small molecule inhibitor to block gap junction communication. Finally, live imaging in the presence of a calcium sensitive dye (Fluo-8 AM) was used to better understand differences in calcium transient activity between the cell networks after 28 days of culture in osteogenic or growth media within the strain-stiffening bottlebrush polymer hydrogel. These results, together with gene expression data suggest progress toward meeting the challenge of developing a three-dimensional cell culture platform that supports the formation and function of human osteocyte-like cell networks *in vitro*.

## Results

### Bottlebrush Polymer Hydrogels Can Mimic Collagen Type-1 Hydrogel Mechanical Properties

Bottlebrush polymer dithiol macromers were synthesized by polymerizing poly(ethylene glycol) methyl ether acrylate (PEGA) via reversible addition fragmentation chain transfer polymerization using a difunctional chain transfer agent (CTA) (Figure S1, Table S1). The CTA trithiocarbonate end-groups were cleaved to free thiols post-polymerization using aminolysis (Figure S2, Scheme S1). These bottlebrush polymer dithiol macromers were then combined with a 20 kDa 8-arm star PEG norbornene in PBS with 0.1 mM CRGDS and a photoinitiator, LAP (6.8 mM), before irradiating with 365 nm light (5 mW/cm^2^) for 5 minutes to covalently crosslink the bottlebrush polymer (BB) hydrogel by thiol-ene photo-click reactions ^19^ (Scheme S2, Figure S3A). Additionally, the collagen type-1 hydrogels (Col1) were formed by diluting the acidic collagen type-1 in 10x PBS before neutralizing the pH to 7 with 1N NaOH. After neutralization, the collagen solution was incubated for 30 minutes at 37 °C to allow the collagen fibrils to form (Figure S3B). Both a nanoporous BB (15 wt%) and fibrillar Col1 (3 mg/mL) hydrogel were formulated to achieve matched *in situ* shear moduli of ∼70 Pa (Figure 1A, Table 1). Notably, the more viscoelastic hydrogel, Col1, exhibited a tan δ of 0.125 whereas the tan δ of the covalently crosslinked BB hydrogel was an order of magnitude lower (0.0125) (Figure 1B, Table 1). As a result, the BB hydrogel exhibited minimal stress relaxation as compared to the Col1 hydrogel that dissipated 50% of applied stress within 100 s (Figure 1D, Table 1). However, both samples showed an onset of stiffening behavior at critical stresses (*σ*_c_) < 25 Pa, within the biologically relevant stress regime (BRSR) (Figure 1C, Table 1). Both hydrogels were measured to have similar stiffening indexes, *m,* of 0.67 for Col1 ^16^ and a slightly less reactive *m* of 0.41 for the BB hydrogel (Table 1). The mechanical properties of both hydrogels are dynamic from *in situ* formulation to *equilibrium swollen* states and in response to cell-matrix interactions. Based on the swollen modulus of the BB hydrogel (40 Pa), we predict that the critical stress experienced by the cells in culture is ∼7.5 Pa ^19^. Conversely, the Col1 hydrogel is known to compact and degrade in the presence of cells^20^, leading to a dynamic critical stress and stress relaxation behavior that is less predictable. Using these two material formulations with similar initial moduli, we next compared the influence of strain-stiffening mechanical stimuli on human mesenchymal stem cells (hMSCs) as a function of stress-relaxation behavior.

**Figure 1.**
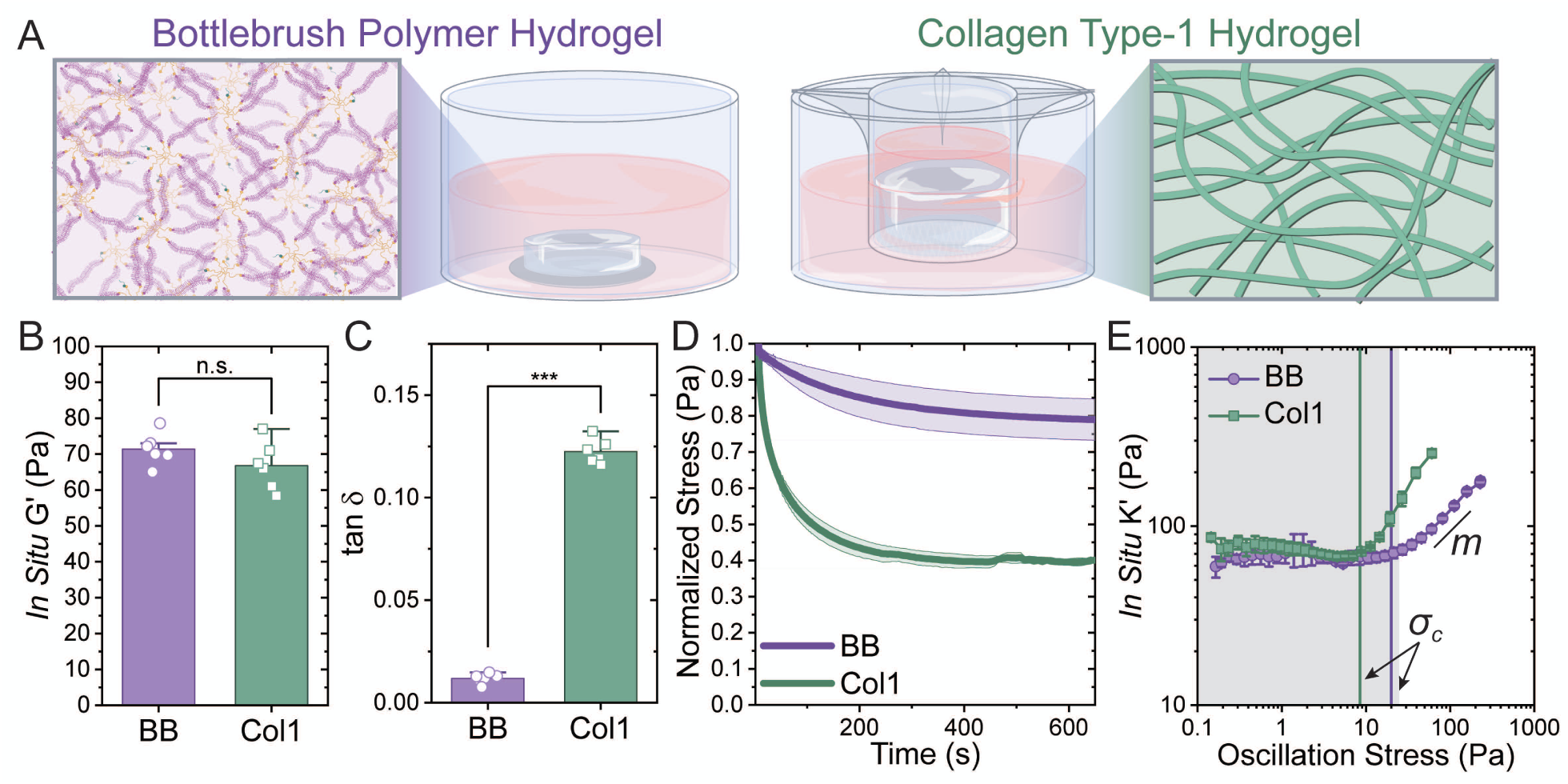
Bottlebrush Polymer Hydrogels Can Mimic Collagen Type-1 Hydrogel Mechanical Properties: Mechanical properties of bottlebrush polymer hydrogels and collagen. A) llustration of network architecture and 3D cell culture set-up for the bottlebrush (BB) polymer hydrogels. BB hydrogels were affixed to thiolated coverslips, and the collagen type 1 hydrogels were cultured in a trans-well insert to reduce rapid compaction. B) *In situ* shear moduli measurements for the BB and Col1 showing both to be ∼70 Pa. C) Plotted tan δ values for the BB and Col1 hydrogels. D) Normalized stress-relaxation after application of 10% strain for the BB and Col1 hydrogels. E) *In situ* differential modulus, K’ plotted as a function of oscillation stress. The critical stress, σ_c_, is labeled at the change from linear to non-linear behavior and the stiffening, *m*, is the slope of *K*’ after σ_c_. The grey box denotes the biologically relevant stress regime (BRSR). (*** *p* < 0.001)

**Table 1.**
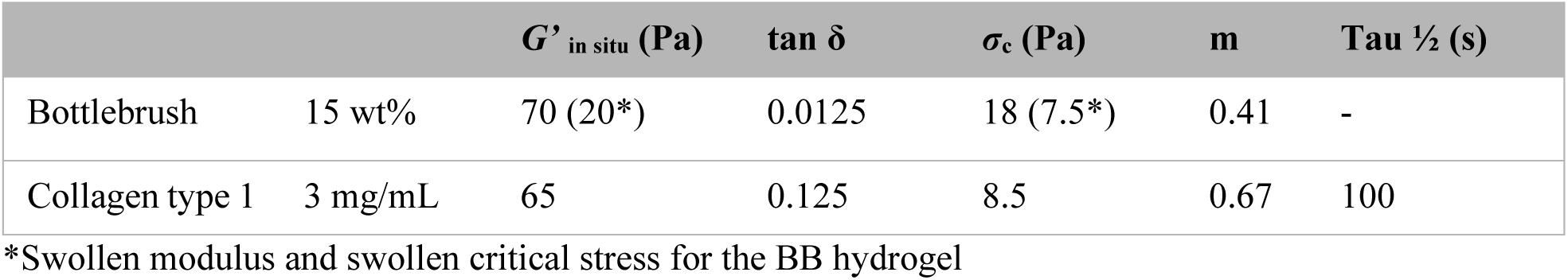
Comparison of hydrogel mechanical properties.

| | | $G'_{\text{in situ}}$ (Pa) | $\tan \delta$ | $\sigma_c$ (Pa) | m | Tau $\frac{1}{2}$ (s) |
| --- | --- | --- | --- | --- | --- | --- |
| Bottlebrush | 15 wt% | 70 (20*) | 0.0125 | 18 (7.5*) | 0.41 | - |
| Collagen type 1 | 3 mg/mL | 65 | 0.125 | 8.5 | 0.67 | 100 |

\*Swollen modulus and swollen critical stress for the BB hydrogel

### Strain-stiffening Dominant Hydrogels Promote Cell Network Formation

hMSCs were encapsulated within 20 μL bottlebrush polymer hydrogels (2.5 million cells/mL), which were formed in cylindrical molds (diameter = 5 mm, height = 1 mm) to adhere one side of the hydrogel to a thiolated coverslip (Figure 1A). For comparison, hMSCs were also encapsulated in Col1 hydrogels. To slow and reduce the effects of cellular compaction in the Col1 hydrogels, hMSCs were seeded at a lower density of 500,000 cell/mL and the initial hydrogel volume was increased to 1 mL to fill a trans-well insert which provided a stronger attachment of the hydrogel to the transwell membrane than standard tissue culture treated multi-well plates and facilitated nutrient diffusion through the thicker hydrogel (Figure 1A). Regardless, the initial total number of cells encapsulated within each scaffold (BB or Col1) was kept similar at 500,000 cells per hydrogel. The reduced encapsulation cell density and increased hydrogel volume in the Col1 systems was used to slow the rate of gel compaction, to better compare the Col1 hydrogels to the BB PEG-based hydrogels where we expect minimal cell-based degradation and contraction of the scaffold dimensions, which would change the cell density (Figure 1A). Cell-laden gels were cultured for up to 28 days in either growth media (GM) or osteogenic media (OM). Live cell imaging indicated high cell viability after 28 days of culture (Figure S4). The hMSC laden gels were also imaged via laser scanning confocal microscopy where the cells appeared to form connections between the actin cytoskeleton in all conditions. As expected, the density of cells within the BB hydrogels remained relatively constant at approximately 2.0 ×10^−6^ nuclei/μm^3^ or 2 million cells/mL (similar to the initial encapsulation concentration). The invariant cell nuclear density in the BB OM is indicative of minimal proliferation and suggestive of differentiation compared to the BB GM condition, which formed multi-nucleated connected cell structures (Figure 2A, B, E). Specifically, with proliferation in the BB hydrogels in GM, the nuclear density increased to 4.0 ×10^−6^ nuclei/μm^3^ (4 million cells/mL) by day 28. Further, the morphology of hMSCs in the BB GM condition was elongated and cellular networks formed with thick, cytoplasm-rich connections (3-9 μm) (Figure 2Av and G). In contrast, hMSCs embedded in BB gels and cultured in OM adopted thinner spindle-like morphologies (2-4 μm) (Figure 2Bv and G) and formed connections with surrounding cells through actin-rich protrusions. In addition, many of the cells in the BB microenvironments appeared to be connected to one another, increasing steadily in the OM and reaching ∼85% connectivity after 28 days (Figure 2F). In GM, hMSCs in the BB hydrogels reached 85% connectivity more quickly, by day 14, and maintained high connectivity for the remaining 14 days of culture (Figure 2F).

**Figure 2.**
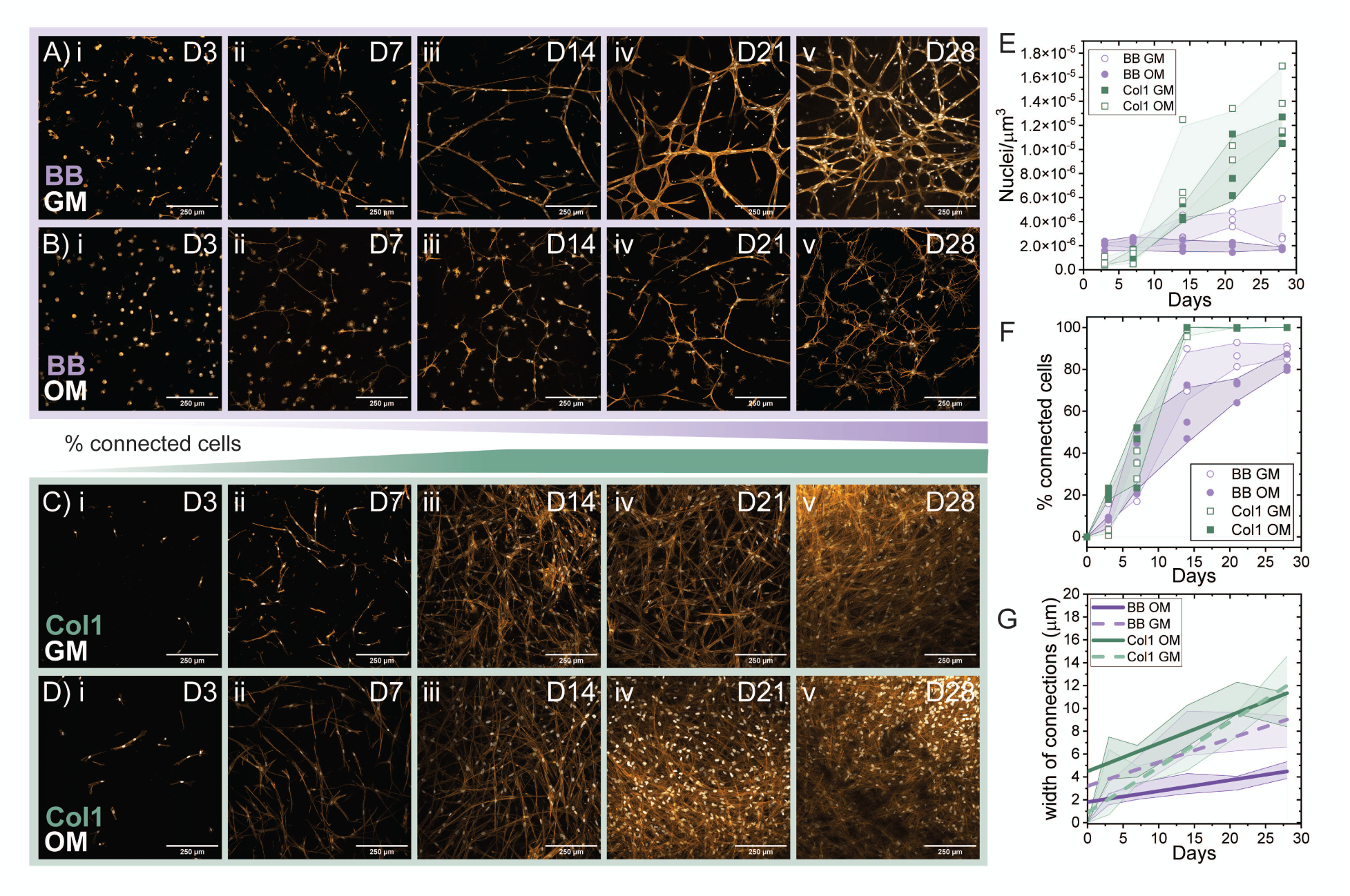
Strain-stiffening Dominant Hydrogels Promote Cell Network Formation: Representative maximum intensity projection (50 μm z-stacks) reveal how cell morphology changes over time [i) D3, ii) D7, iii) D14, iv) D21, and v) D28] for hMSCs cultured in the bottlebrush polymer scaffold and GM (A) or OM (B) and the collagen type-1 scaffold and GM (C) or OM (D). F-actin was visualized using rhodamine phalloidin (orange) and DNA with DAPI (white). The cell network evolution was quantified as a function of time and defined by E) the number of nuclei per μm^3^, F) the percentage of visually connected cells in the network, and G) the width of the f-actin connections between nuclei.

Of note, the minimally stress relaxing BB hydrogels led to notably different hMSC morphologies and network organizations compared to that observed in the Col1 hydrogels. In the Col1 hydrogels, the density of the cells, as quantified by the nuclei per imaging volume, increased dramatically over the first week of culture. By day 14 of culture, it increased from 0.4 ×10^−6^ nuclei/μm^3^ (approximately 400,000 cells/mL) at day 3 to greater than 4.0 ×10^−6^ nuclei/μm^3^ (4 million cells/mL), presumably due to cell-mediated degradation of the Col1 matrix and/or its compaction (Figure 2C-E). At day 28, the average nuclear density in the Col1 hydrogels was 11 ×10^−6^ and 14 ×10^−6^ nuclei/μm^3^ (11 million and 14 million cells/mL) for the OM and GM, respectively (Figure 2E and Figure S5). This dramatic increase in nuclear density over the course of 14 days led to >95% of connected hMSC in the Col1 matrix across both culture conditions up to 28 days (Figure 2F). Further, the width of actin connections between nuclei was similar to actin connections in the BB GM condition and increased from ∼5 μm to > 8 μm by day 28 for the OM condition and ∼ 12 μm in diameter for the GM condition (Figure 2G).

### Osteogenic Media Promotes Connection-Dependent Fluorescence Recovery

Given the pronounced compaction observed in the Col1 hydrogels (Figures 2C-D and S5) and the marked differences in cell morphology and network architecture across hydrogel and media conditions, we next sought to determine whether these morphological differences translated into functional differences in intercellular connectivity. To characterize this, we performed fluorescence recovery after photobleaching (FRAP) experiments on live cell networks after 28 days of culture to assess functional connectivity between neighboring cells. Prior to imaging, all samples were incubated for 30 minutes with Calcein AM (0.5 μM) and extensively washed with PBS to remove exogenous dye from the hydrogel, ensuring that fluorescence recovery reflected intracellular transport rather than extracellular diffusion. A region of interest (ROI) was drawn around a cell of interest within each of the samples and the fluorescence intensity within the ROI was monitored as a function of time. Figure 3A-D shows representative z-stack sum slice images to characterize the fluorescence intensity 30 seconds prior to photobleaching (−00:30), immediately after the light exposure within the ROI (00:00), and 9 minutes and 30 seconds after the initial photobleaching (9:30). Representative videos of FRAP experiments for all four samples, including an isolated cell control, are presented in the Supplementary Information (Videos S1-5). The lack of compaction in the bottlebrush hydrogels and 85% connectivity allowed identification of isolated cells within the network, which served as internal negative controls to confirm that fluorescence recovery arose from intercellular connections rather than residual dye uptake (Figure 3E, S6). These experiments enabled direct comparison of connection-dependent dye dynamics between the two hydrogel platforms and in the presence or absence of differentiation conditions.

**Figure 3.**
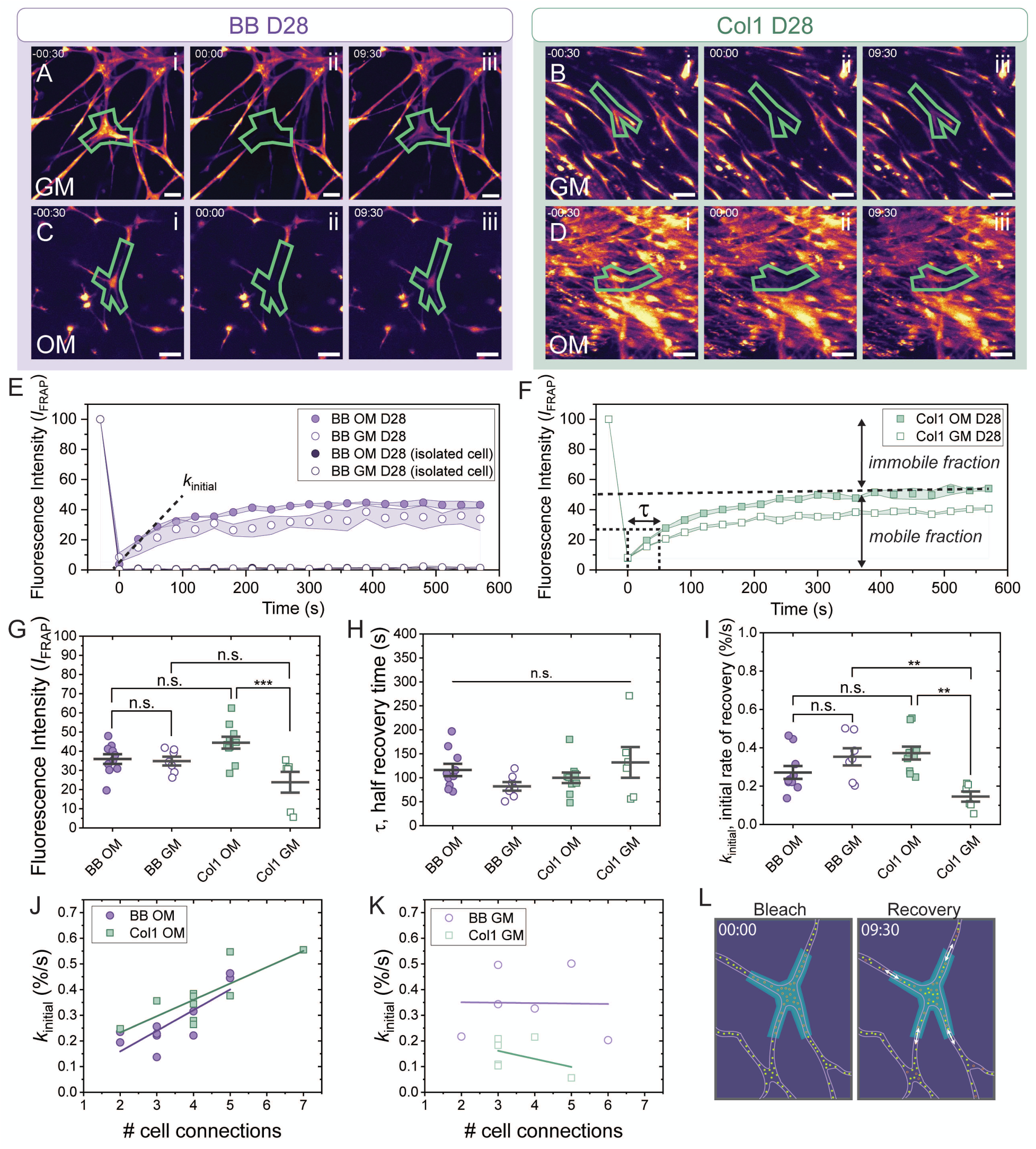
Osteogenic Media Promotes Connection-Dependent Fluorescence Recovery: Fluorescence recovery after photobleaching experiments to confirm cell network connectivity. Representative sum-slice z-projections (scale bar = 50 μm) for A) BB GM, B) Col1 GM, C) BB OM, and D) Col1 OM after 28 days of culture showing the cell within the region of interest (green) prior to bleaching (i), immediately after bleaching (ii), and after 9 minutes and 30 seconds of recovery (iii). Quantification of the normalized fluorescence intensity, *I*_FRAP_, over time for samples cultured in the E) BB hydrogels and F) Col1 hydrogels. Comparisons between samples of the G) total *I*_FRAP_ recovery (representative of the mobile fraction), H) half recovery time, τ, and I) initial rate of recovery, *k*_initial_. (n = 2-5 ROI per hydrogel across 3 biological replicates, \**P*<0.05, \*\**P*<0.01, \*\*\**P*<0.001) where *k*_initial_ is plotted as a function of the number of visible connecting cells plotted for J) OM and K) GM conditions. L) Illustration of the photo-bleaching and fluorescence recovery process of Calcein AM molecules within the cell of interest.

For each of the hMSC-laden hydrogel conditions, with the exception of isolated cells (Figure S6), effective photobleaching and subsequent fluorescence recovery were observed over the course of 10 min. Recovery curves were fit using the following model:

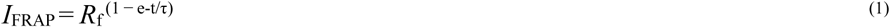

Here, fluorescence intensity (*I*_FRAP_) is defined as a function of the mobile or recovery fraction (*R*_f_) that represents the plateau in fluorescence recovery, and τ denotes the characteristic recovery time. Relatively modest differences in *R*_f_ and τ (Figure 3G–H) were observed between conditions, where each BB polymer had a similar amount of fluorescent recovery (40%); however, cells in the GM Col1 samples exhibited a lower amount of fluorescent recovery (25%) than those in OM Col1 hydrogels (40%). Similarly, the initial recovery rate (*k*_initial_) over the first 60 s revealed minimal differences in transport dynamics between the BB OM, BB GM, and Col1 OM conditions, all of which achieved similar initial rates of recovery (approx. 0.35 %/s). However, cells within the Col1 GM condition recovered more slowly with a *k*_initial_ of < 0.2 %/s (Figure 3I).

More notably, when the initial recovery rate was plotted as a function of the number of visible connections to neighboring cells, distinct trends emerged depending on the media formulations (Figure 3J– K, Table S2). In GM, there was little dependence of the recovery rate on the number of cell-cell connections (Pearson’s R < −0.45). In contrast, in OM, a strong positive linear correlation (Pearson’s R ≥ 0.80) was observed between the initial recovery rate and the number of intercellular connections, indicating that functional connectivity increased with network complexity and differentiation (Table S2). Based on the substantially larger width of the intercellular connections observed in the BB GM condition relative to the BB OM (Figure 2G), we hypothesized that distinct mechanisms may be underlying network formation in these two culture conditions. Specifically, cells in GM may undergo partial cell fusion or achieve cytoplasmic continuity, whereas cells in OM may preferentially form thinner, dendrite-like connections.

### Gap Junctions Exist Along Connected Cell Protrusions

To further interrogate potential mechanisms underlying functional connectivity within the developing cell networks, we examined the presence of connexin-43 (Cx43), a transmembrane protein expressed by osteoblasts and osteocytes during bone development and remodeling (Figure 4A-D). Cx43 forms hexamers along the cell membrane (i.e., hemichannels) that facilitate the exchange of signaling factors between the cell cytoplasm and extracellular microenvironment. When two hemichannels align along the cell membranes of adjacent cells, they form a gap junction that facilitates intercellular communication^21,22^ (Figure 4H). Immunostaining for Cx43 revealed a greater number of puncta associated with the f-actin staining from cells cultured in GM than in OM across both hydrogel systems (Figure 4A-D). This trend suggests that expression of Cx43 alone is not sufficient to explain the distinct functional connectivity observed by FRAP, particularly given the enhanced connection-dependent recovery observed in osteogenic conditions.

**Figure 4.**
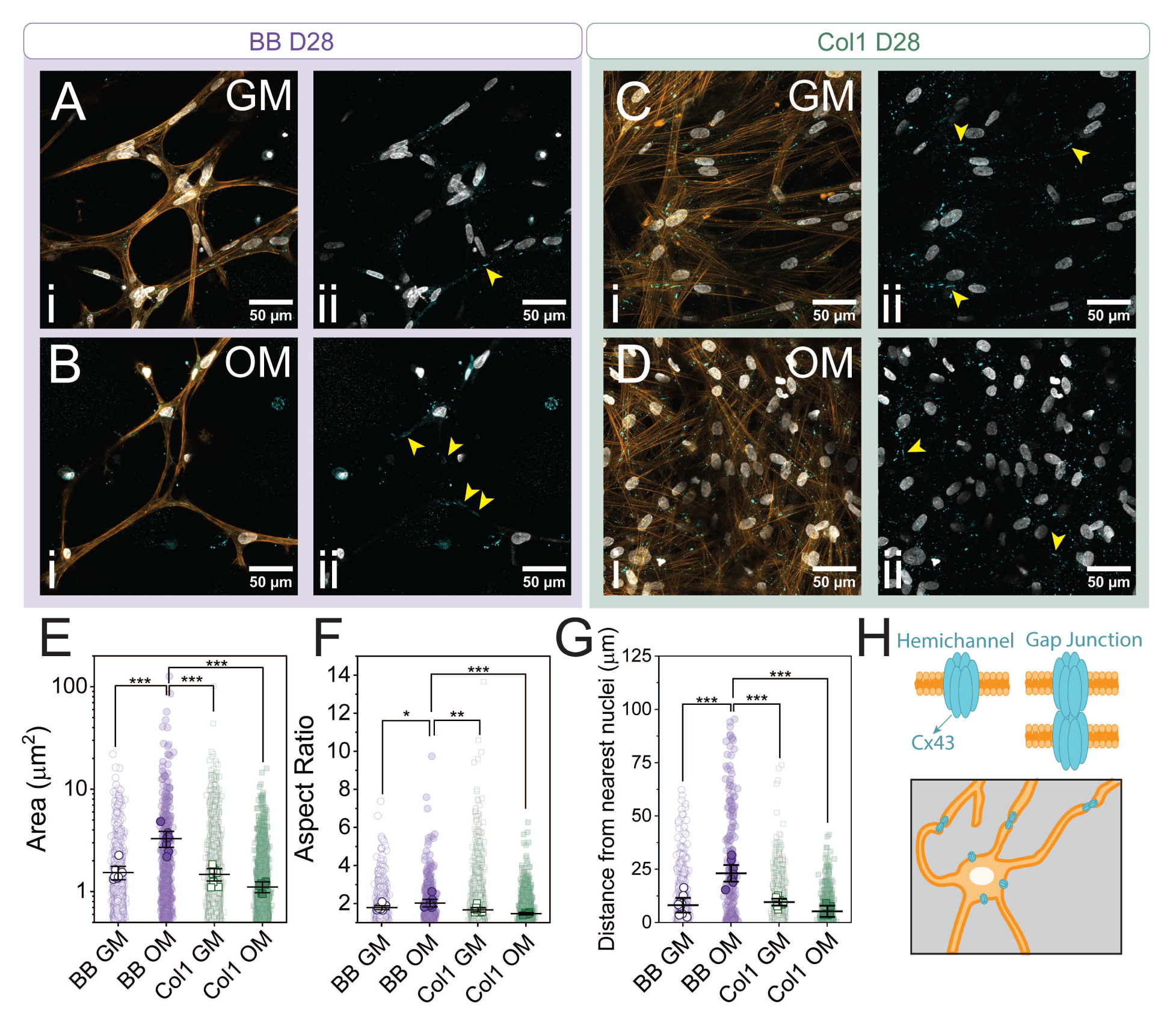
Gap Junctions Exist Along Connected Cell Protrusions: Connexin 43 immunocytochemical staining across all four culture conditions on day 28. Representative maximum intensity projections of confocal images for A) BB GM, B) BB OM, C) Col1 OM, D) Col1 OM where i) is an overlay of actin (orange), DNA (white), and connexin 43 (cyan) and ii) is an overlay of DNA (white) and connexin 43 (cyan). Yellow arrows indicated locations of high aspect ratio connexin 43 puncta. Quantification of the E) area, F) aspect ratio, and G) distance from nearest nuclei of connexin 43 puncta. (n = at least three fields of view across 3 biological replicates, \**P*<0.05, \*\**P*<0.01, \*\*\**P*<0.001) H) illustration of connexin 43 hexamers forming hemichannels or gap junctions.

When visualized through immunocytochemical staining, the Cx43 proteins were bright puncta along the actin cytoskeleton (Figure 4A-D). This Cx43 staining can either be indicative of hemichannels on the cell membrane, which are involved in the influx and efflux of biochemical signals between the cell and surrounding matrix in response to mechanical stimulus, or as gap junctions that bridge two cell membranes and facilitate the transport of ions and nutrients (Figure 4C-F). To differentiate between gap junctions and hemichannels, the area, aspect ratio, and distance from the nearest nuclei of the Cx43 puncta were quantified (Figure 4E-G). When Cx43 was stained in primary osteoblasts and osteocytes cultured in 2D, the Cx43 distribution in osteoblasts presents as scattered puncta throughout the cell cytoplasm near the nucleus and along cell-cell contacts. In primary osteocytes, the Cx43 puncta are also observed in the cytoplasm, but the majority are localized to the intersection of dendritic processes^23^. After 28 days of culture, the area of the Cx43 puncta observed in cells within the BB OM condition was significantly larger than the three other conditions (Figure 4E). In addition, the average aspect ratio of the Cx43 puncta observed on the cytoskeleton cells cultured in the BB OM condition was larger than the other three conditions (Figure 4F). Interestingly, the average aspect ratio was the smallest for the Cx43 puncta quantified in the Col1 OM condition. Further, we observed that many of the Cx43 puncta in the BB OM condition were along the dendrite-like protrusions of the cell network, so the distance between each Cx43 puncta and the nearest nuclei was quantified. This analysis confirmed that the Cx43 puncta in the BB OM condition were, on average, significantly further away from the cell nuclei than the other three conditions, suggesting localization to the protrusions (Figure 4G). Together, the larger area and aspect ratio of Cx43 puncta that are located further away from cell nuclei in the BB OM condition are more indicative of gap junctions along the cell protrusions than hemichannels on the cell membrane (Figure 4Bii).

### Connexin-mediated Transport Facilitates Intercellular Communication

Next, to assess the role of Cx43-mediated communication in functional network connectivity, we performed live-cell FRAP experiments at day 28 while pharmacologically inhibiting gap junctions with carbenoxolone (CBX). hMSCs were treated with 100 µM CBX for 24 h (Figure S7)^24^ prior to FRAP measurements, while CBX was maintained in the imaging media to ensure continuous blockade of connexin hemichannels and intercellular gap junctions during data acquisition (Figure 5A-D, Videos S6-9). For all conditions, CBX treatment resulted in a measurable reduction in both the total fluorescence recovery and the initial rate of recovery (Figure 5E-G), confirming that connexin-mediated transport contributes to intercellular communication within these networks.

**Figure 5.**
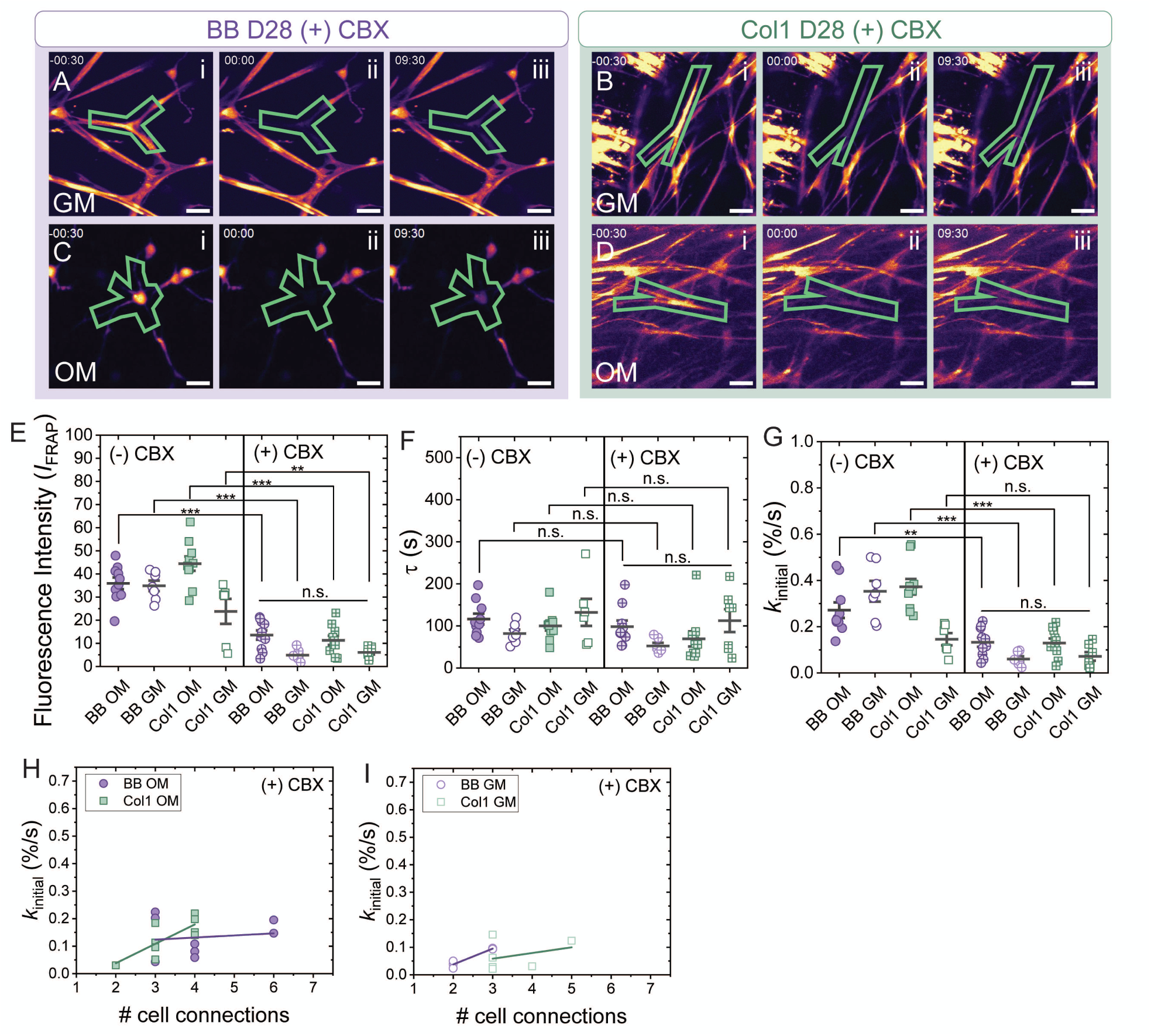
Connexin-mediated Transport Facilitates to Intercellular Communication: Fluorescence recovery after photobleaching experiments after 24 hour treatment with 100 μm CBX. Representative sum-slice z-projections (scale bar = 50 μm) for A) BB GM, B) Col1 GM, C) BB OM, and D) Col1 OM (scale bar – 50 μm) after 28 days of culture showing the cell within the region of interest (green) prior to bleaching (i), immediately after bleaching (ii), and after 9 minutes and 30 seconds of recovery (iii). Comparisons between untreated, (−) CBX, and treated, (+) CBX, showing the overall decrease in E) total *I*_FRAP_ recovery (representative of the mobile fraction), F) the half recovery time, τ, and G) the initial rate of recovery, *k*_initial_ after treatment with CBX. (n = 2-5 ROI per hydrogel across 3 biological replicates, \**P*<0.05, \*\**P*<0.01, \*\*\**P*<0.001). *k*_initial_ is plotted as a function of the number of visible connecting cells for H) OM and I) GM conditions.

To further dissect these effects, we examined how CBX treatment altered the relationship between functional recovery and network architecture (Figure 5H-I). In the absence of CBX, the BB OM condition exhibited a strong positive linear correlation (Table S2, Pearson’s R = 0.82) between the initial FRAP recovery rate and the number of visible connections to neighboring cells. Strikingly, CBX treatment eliminated this correlation in the BB OM condition (Pearson’s R = 0.14), indicating that Cx43-mediated gap junctions are the dominant pathway supporting connection-dependent communication in this system. In contrast, the Col1 OM condition retained a positive correlation (Pearson’s R = 0.78) between recovery rate and cell connectivity even in the presence of CBX, suggesting that alternative avenues of intercellular communication persist within the compacted collagen matrices. Notably, in both GM conditions, there was low fluorescence recovery (< 10%) suggesting minimal intercellular communication in the presence of CBX (Figure 5E).

### Strain-stiffening Dominant Bottlebrush Polymer Hydrogels Promote Early Osteocyte Gene Expression

As further evidence of the formation of a functional osteocyte-like network, we measured the expression of genes associated with an osteoid osteocyte phenotype, osteopontin (OPN) and E11, also known as podoplanin, (E11/PDPN) across the 28-day differentiation timeline and for each of the four culture conditions. Fold change expression was normalized to hMSCs at D0 (prior to encapsulation in the hydrogel scaffolds). We observed a significant fold change increase in OPN expression starting on day 14 following a significant increase in OCN expression at day 3 in the BB OM condition (Figure 6A, Figure S9D), as well as the early osteocyte marker, E11/PDPN (Figure 6B). E11 is linked to protrusion extension and dendritic network formation in osteoid-osteocytes ^25^. Interestingly, at day 28 of culture in the BB GM condition, we also observed a fold change increase in E11/PDPN expression, which we attributed to the cell network formation that was observed in Figure 2A. We posit that regardless of the osteogenic biochemical signals present in the media, hydrogel scaffolds that are strain-stiffening dominant over stress-relaxation upregulate E11 expression, and this is further tied to cellular morphological changes and network formation descriptive of osteoid osteocytes. Contrastingly, we observed minimal changes in OPN or E11 expression over the 28-day culture period from cells isolated from Col1 scaffolds cultured either in GM or OM.

**Figure 6.**
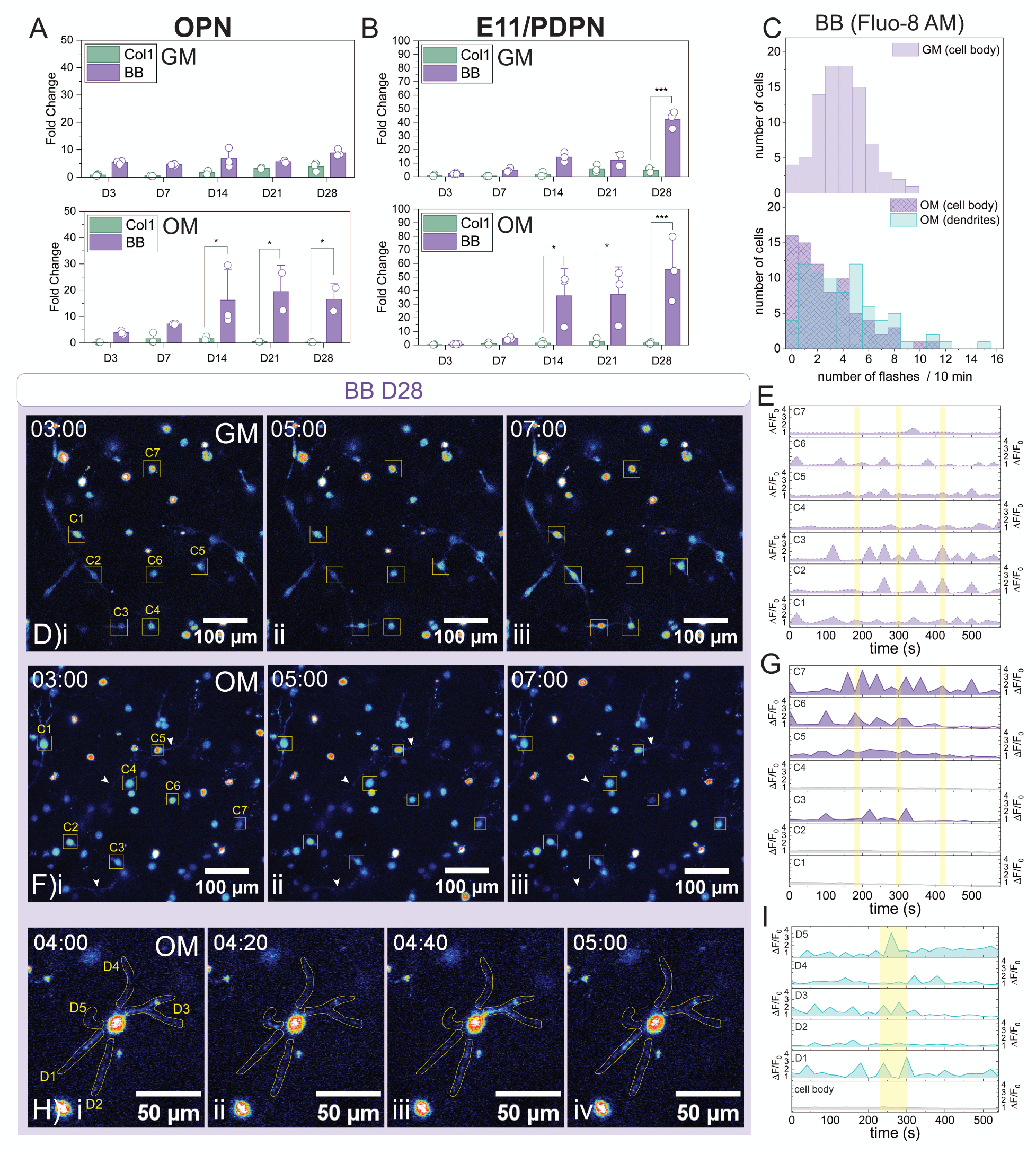
Osteoid osteocyte-like cell gene expression and cell network calcium transient activity. qPCR fold change in RNA expression normalized to D0 (day of encapsulation) measured at D3, D7, D14, D21, and D28 in both the BB and Col1 hydrogels in GM and OM for the osteogenic marker A) OPN and early osteoid-osteocyte marker, B) E11/PDPN. (n = 2-3 biological replicates (containing RNA from 3-5 pooled hydrogels at each timepoint), \**P*<0.05, \*\**P*<0.01, \*\*\**P*<0.001) Calcium transient events visualized using Fluo-8-AM in the BB hydrogels after 28 days of culture in GM or OM summarized as C) the number of cell bodies or cell dendrites which flash a defined number of times over the course of 10 minutes. Representative images of sum-slice z-projections of the Fluo-8 AM fluorescence intensity in the D) BB GM and F) BB OM at i) 3 minutes, ii) 5 minutes, and iii) 7 minutes for seven representative cells of interest for which the change in normalized change in fluorescence intensity (ΔF/F_0_) is plotted over 10 minutes in E) and G), respectively. Yellow bars indicate the quantification of the images at 3, 5, and 7 minutes. H) Representative Fluo-8 AM sum-slices z-projection image series show changes in normalized fluorescence intensity (ΔF/F_0_) for cellular dendritic protrusions at i) 4 minutes, ii) 4 minutes and 20 seconds, iii) 4 minutes and 40 seconds, and iv) 5 minutes. I) Quantification of the change in normalized fluorescence intensity (ΔF/F_0_) over 10 minutes for 5 representative dendrites. The yellow region corresponds to the image series shown in H.

### Osteoid Osteocyte-like Cell Dendrites Contribute to Mechanosensing

Given the observed increase in expression of the osteoid osteocyte marker, E11, in the BB hydrogel culture conditions, we next investigated the calcium transient behavior for these samples in GM and OM after 28 days of culture. Un-differentiated hMSCs are known to exhibit spontaneous increases in intercellular calcium without external stimulation^26^ that often corresponds to release of intercellular calcium stores from the endoplasmic reticulum^27^. However, osteocytes are key mechanosensors in bone ^28^ and will also experience an influx in intercellular calcium in response to activation of mechanosensitive ion channels. To investigate any differences in the calcium transients between the GM and OM condition, we incubated cells in the BB hydrogels with a fluorescent calcium sensitive dye, Fluo-8 AM, prior to imaging for 10 minutes on a laser scanning confocal microscope. In GM, we observed that a majority of cells flashed 2-6 times over the course of 10 minutes (Figure 6C). These flashes were spontaneous and occurred across the cell body and visible connection points between cells. Comparatively, ∼30% of cells cultured in OM either did not experience an influx in intercellular calcium or experienced only one calcium transient event over 10 minutes (Figure 6C). Because we applied no external chemical agonists or mechanical stimuli to the samples during imaging, the low percentage of cells with minimal calcium ion influx events was not surprising. However, when changes in the normalized fluorescence intensity (ΔF/F_0_) for the Fluo-8 AM signal was plotted as a function of time (Figure 6D-G), we saw that spikes of varied fluorescence intensity occurred sporadically over the course of the 10 minutes. In contrast, cells in OM within the cellular network (i.e., C1, C2, and C4) did not experience temporal changes in intercellular calcium concentration over the duration of the measurement (Figure 6F-G). Upon further investigation, although the cell bodies did not flash, we observed that the cellular dendrites experienced independent changes in calcium concentration over the course of 10 minutes (Figure 6C, H-I). In fact, the majority of cell dendrites underwent one to five flashes over the course of 10 minutes (Figure 6C). As seen in Figure 6H, while the cell body did not have a measurable change in fluorescence intensity, each of the dendrites (D1-5) underwent independent increases in fluorescence intensity (localized calcium concentration) with D1 and D3 experiencing the most spikes. These results corroborate the hypothesis that osteocyte dendrites can be mechanically sensitive and that they can respond independently to mechanical stimulus compared to the osteocyte like cell bodies^29^. This suggests that dendrites could be experiencing membrane tension changes that can open and close ion channels on the protrusions as the three-dimensional cellular network exerts forces on the surrounding osteoid-mimetic synthetic extracellular matrix.

## Discussion

As bone formation through intramembranous ossification and bone remodeling both rely on the support of a non-calcified, collagen type 1 rich osteoid to support osteocytogenesis, we sought to isolate the contribution of two osteoid-like mechanical cues, nonlinear elasticity and stress relaxation, to better understand the role each plays on osteogenic cell network formation. To answer this question, we compared a nonlinear elastic BB polymer hydrogel with a shear modulus-matched to a viscoelastic Col1 hydrogel. Both hydrogel systems exhibited critical stresses within the BRSR (< 25 Pa) where fibrillar ECM components, such as collagen and fibrin, undergo stiffening due to cell generated forces^30,31^. However, only the Col1 hydrogel underwent notable stress-relaxation that led to rapid, cell-mediated compaction over the course of the 28-day culture period (Figure 2C-G). Beyond 21 days of culture in the Col1 hydrogels, increases in cell density made imaging deeply into the hydrogel sample difficult due to light attenuation, notable by the high nuclear density and nearly 100% connectivity by day 14 in both GM and OM culture conditions. In contrast, the BB hydrogel maintained a relatively constant cell density in both GM and OM conditions, which enabled direct observation of cellular network formation through the extension of actin-rich protrusions with minimal matrix compaction (Figure 2A-B, E-G). Each of the BB polymer images were collected away from the surface, at least 50 µm into the hydrogel, which made identifying and tracking individual cells and their connections points much easier than in Col1 samples. In general, the BB hydrogel system with a lower tan δ provided a robust platform that supported multi-week cell culture, maintaining consistent cell densities while permitting analysis of morphological changes and osteocyte-like cell network formation.

By day 28, live cell FRAP experiments revealed that intercellular transport scaled positively with the number of visible connections to neighboring cells in OM (Figure 3J), indicating that increased network complexity translated directly into enhanced communication. In contrast, fluorescence recovery in GM conditions showed little dependence on connection number (Figure 3K), which was consistent with broader cytoplasmic signaling rather than transport via gap junctions. These functional differences were not apparent from morphology measurements alone and reinforced the importance of assessing network function, in addition to structure, when modeling osteogenic differentiation. In addition, while Cx43 puncta were observed at day 28 of culture in all four samples (Figure 4A-D), how Cx43 was functionally deployed depended on the matrix architecture and cellular differentiation state. Cx43 can exist on cell membranes as hemichannels or assemble into gap junctions at sites of direct cell–cell contact^23^. We posited that in conditions with high overall Cx43 staining, such as the Col1 hydrogels and BB GM samples, a substantial fraction of Cx43 were hemichannels that contribute to nonspecific molecular exchange (Figure 4A, C, and D). By contrast, the higher aspect ratio of the Cx43 puncta observed in the BB OM condition was more consistent with localized gap junction assembly at discrete contact points (Figure 4B), aligning with the strong dependence of functional recovery on visible cell connections and the pronounced sensitivity to CBX treatment (Figure 5H)^13^.

Taken together, these results suggest that differentiation-dependent remodeling of cell–cell interfaces, rather than total Cx43 expression alone, governs the mode of intercellular communication within developing osteogenic cell networks. As a result, measurement of osteogenic markers, such as osteopontin (OPN) and E11/PDPN, implicated the transition from osteoblast to early embedding osteocyte were more expressed by cells embedded within BB hydrogels compared to Col1 gels and independent of media condition (Figure 6A-B). Since E11 is indicative of cell protrusion extension during osteocytogenesis^32^ its elevated expression highlights how nonlinear elastic biomechanical cues can influence dramatic phenotypic changes compared to cell-dictated viscoelastic ECM remodeling of a non-calcified osteoid. Moreover, these results underscore how matrix mechanics and stability, particularly reduced stress-relaxation and limited compaction afforded by bottlebrush hydrogels, can enable elucidation of connection-specific communication mechanisms that are otherwise obscured in dynamically compacting collagen matrices.

Finally, after extended culture time, hMSCs cultured in GM within the BB hydrogels still exhibited spontaneous, non-coordinated changes in intracellular calcium concentrations (Figure 6E). In contrast after 28 days, cells cultured in OM in the BB osteoid-mimetic hydrogels had lower spontaneous calcium concentration changes, dramatic enough to cause changes in the overall cell body fluorescence intensity. It is worthwhile noting that no mechanical stimulus or chemical agonist were applied during these measurements, so we hypothesize that the cellular networks may not be experiencing a sufficient stimulus to elicit a total cell body calcium transient event. However, the connected cell dendrites underwent local changes in calcium concentration (Figure 6G). The isolated calcium transient events along the cell dendrites could be facilitated by shear flow sensitive ion channels which are known to exist along osteocyte dendrites within the lacunar-canalicular matrix^28,33^. We posit that the dendrites are responding to localized micro-scale poroelastic events inducing small amounts of shear flow or osmotic pressure changes as the cells pull on the nanoporous matrix. In MLO-Y4 cells, hypotonic stimuli applied at dendrite tips similarly induce an increase in calcium concentration within dendrites of interest^34^. Thus, with the ability to image human-derived osteocyte-like cell networks in BB hydrogels in three dimensions, future research directions might investigate which ion channels could drive this change in intercellular calcium signaling. In addition, our prior work demonstrates that hMSC protrusions formed within BB hydrogels can exert forces and displacements within these hydrogel over long distances (> 50 μm)^19^. It would be interesting to model fluid flow and micro-scale poroelastic behavior within the synthetic hydrogel scaffold that result from these cell-induced displacements. Of further note, the osteoid-mimetic synthetic extracellular matrices developed in this study have the potential to enable and validate a human osteocyte 3D *in vitro* model that would be valuable for investigating how changes in cellular mechanotrasduction and cell-ECM signaling influence intramembranous ossification and bone remodeling.

In conclusion, nanoporous synthetic BB hydrogels exhibit strain-stiffening behavior under applied stress while minimizing stress relaxation. The BB hydrogels presented in this study, support multi-week 3D culture of hMSCs while circumventing issues of progressive compaction and concomitant high cell density that occur when using Col1 hydrogels. While maintaining a relatively constant cell density, ,BB matrices enabled hMSCs to undergo pronounced morphological remodeling, extending long cellular processes from a central cell body prior to either fusing with neighboring cells to form multinucleated networks in GM or organizing spindle-like, actin-rich protrusions with functional gap-junction-mediated connectivity in OM. Notably, the latter cell network architecture closely resembles the dendrite-rich connectivity observed between embedded osteogenic cells within non-mineralized osteoid during intramembranous ossification. These findings suggest that persistent nonlinear elastic cues, in the absence of substantial stress relaxation, are sufficient to promote osteocyte-like network organization independent of matrix compaction. Building on this platform, future studies that incorporate biomimetic and/or biochemical cues into BB hydrogels may enable differentiation and further maturation of hMSCs into functional osteocyte networks, thereby offering new opportunities to investigate osteocyte mechanotransduction and to develop osteocyte-mediated disease models *in vitro*.

## Materials & Methods

### Hydrogel Synthesis Materials

2-(dodecylthiocarbonothioylthio)-2-methylpropionic acid (DDMAT, Sigma-Aldrich), ethylene glycol (Sigma-Aldrich), 4-dimethylaminopyridine (DMAP, Sigma-Aldrich), N,N’-diisopropylcarbodiimide (DIC, Sigma-Aldrich), poly(ethylene glycol) methyl ether acrylate [average MW = 480 Da] (PEGA9, Sigma-Aldrich), 2,2’-Azobis(2-methylpropionitrile) (AIBN, Sigma-Aldrich), N’N-dimethylformamide (DMF, anhydrous, 99.8%, Alfa Aesar), RC Dialysis Tubing (MWCO 3.5 kDa, Spectrum Spectra/Por 3, Fisher Scientific), HCl (2M, Sigma-Aldrich), Tris(2-carboxyethyl)phosphine hydrochloride (TCEP, Sigma-Aldrich), n-butylamine (Sigma-Aldrich), Endo-exo-5-norbornene-2-carboxylic acid (NB-COOH, Sigma-Aldrich), 20 kDa 8-arm poly(ethylene glycol) amine (tripentaerythritol), HCl Salt (JenKem, ≥ 95% substitution), O-(7-Azabenzotriazol-1-yl)-N,N,N’,N’-tetramethyluronium hexafluorophosphate (HATU, ≥ 99.5% (HPLC), Chem-Impex), N,N-Diisopropyletheylamine (DIPEA, Sigma-Aldrich), DPBS (no calcium no magnesium, Fisher-Scientific), Lithium phenyl-2,4,5-trimethylbenzoylphosphinate (LAP, Sigma-Aldrich), CRGDS (Bachem). Rat tail collagen type 1 (Corning), 10x PBS (ThermoFisher), NaOH (1N, Sigma-Aldrich)

### Cell culture and Bioassay Materials

Human mesenchymal stem cells (Rooster Bio, lot: 310305, donor: female, 20 y.o.), RoosterNourish media (RoosterBio), Low-glucose Dulbecco’s Modified Eagle medium (DMEM, ThermoFisher), Phenol-free low glucose DMEM (ThermoFisher), Fetal Bovine Serum (FBS, ThermoFisher), Penicillin (ThermoFisher), Streptomycin (ThermoFisher), Amphotericin B (Fungizone, ThermoFisher), L-ascorbic acid (Sigma-Aldrich), beta-glycerol phosphate (Sigma-Aldrich), dexamethasone (Millipore Sigma), Carbenoxolone (Sigma-Aldrich), Rhodamine Phalloidin (Invitrogen), DAPI (Sigma-Aldrich), Connexin 43 antibody (rabbit) (Abcam, ab314908), Alexaflor 488 Goat anti-rabbit (ThermoFisher), Calcein AM (Invitrogen), Fluo-8-AM (Abcam), Pluronic F-127 (Sigma-Aldrich), Presto Blue (Fisher-Scientific), 10% formalin (Sigma-Aldrich), Tween 20 (Millipore Sigma), Triton X-100 (Fisher-Scientific), Trizol (Invitrogen), chloroform (Sigma-Aldrich), RNeasy micro kit (Qiagen), ethanol (Fisher-Scientific)

PCR Primers

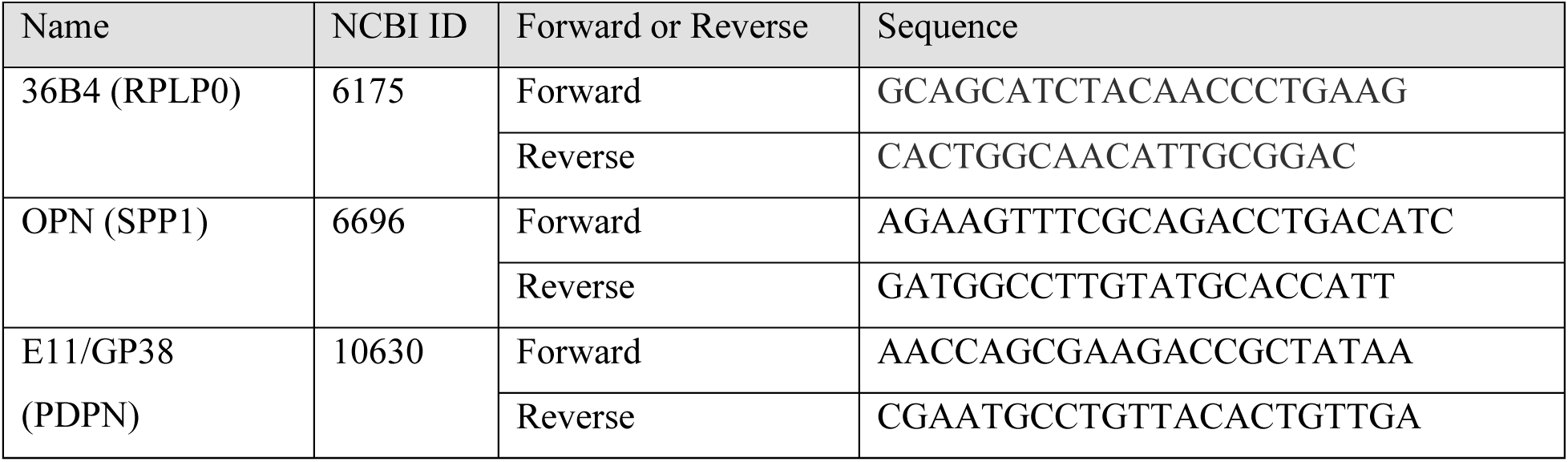

### Cell culture – expansion and differentiation

hMSCs were purchased from RoosterBio and expanded following RoosterBio’s expansion protocols. Prior to encapsulation, hMSCs (passage number = 4) were seeded for expansion on tissue culture treated poly(styrene) at 4,000 cells/cm^2^ and cultured in RoosterNourish media until 70% confluent. The cells were then encapsulated in either bottlebrush polymer or collagen type-1 hydrogel scaffolds using the procedures described below. Growth media conditions were cultured in low-glucose DMEM media supplemented with 10 v/v% FBS, 1 v/v% penicillin-streptomycin, and 0.1 v/v% fungizone. This media was then further supplemented with L-ascorbic acid (50 μM), beta-glycerol phosphate (10 mM), and dexamethasone (0.1 μM) to make osteogenic media. GM and OM was changed every other day.

### Hydrogel Synthesis and Cell Encapsulation

#### Bottlebrush Polymer Dithiol Synthesis

The bottlebrush polymer (C_12_H_25_-PPEGA9-100-C_12_H_25_) was synthesized via reversible addition fragmentation chain transfer polymerization with a difunctional 2-(dodecylthiocarbonothioylthio)-2-methylpropionic acid (di-DDMAT) chain transfer agent and PEGA9. The difunctional chain transfer agent was synthesized via a carbodiimide coupling between two DDMAT molecules and ethylene glycol. Detailed synthetic methods for di-DDMAT are reported previously by Ohnsorg *et al.*^19^. PPEGA9 (550 eq.) was combined with di-DDMAT (1 eq.), and AIBN (0.1 eq.) in anhydrous DMF (1.0 M). The reaction mixture was purged with N_2_ for 30 minutes prior to heating at 70 °C overnight. The reaction mixture was quenched in LN_2_ to end the polymerization, and the polymer was purified by dialysis (8 kDa MWCO, RC Dialysis bag) in methanol. Following dialysis, the polymer was isolated from methanol via rotary evaporation followed by further drying under high vacuum. The polymer was characterized via ^1^H NMR in CDCl_3_ and SEC-MALS (Tosoh EcoSEC) in DMF with 0.1% LiBr (Figure S1 and Table S1).

The trithiocarbonate end-groups were then cleaved to free thiols using aminolysis. The C_12_H_25_-PPEGA9-100-C_12_H_25_ (2 eq.) was dissolved in THF (0.2 M) along with TCEP (1 eq.) before adding n-butylamine (100 eq.) dropwise to the reaction mixture and stirring at room temperature for 2 hours. The light-yellow reaction mixture became colorless after 30 minutes. The polymer was then purified by dialysis (8 kDa MWCO) in methanol. On the second dialysis media change, the polymer was dialyzed against acidic methanol (pH = 3-5) for one media change. After at least four more dialysis media changes, the bottlebrush polymer dithiol (SH-PPEGA9-100-SH) was isolated from the methanol via rotary evaporation followed by further drying under high vacuum. The percentage of trithiocarbonate end groups removed was characterized by ^1^H NMR in CDCl_3_ (Schematic S1 and Figure S2).

#### Bottlebrush Polymer Hydrogel Synthesis

The bottlebrush polymer hydrogels (15 wt%) were crosslinked using a photoinitiated thiol-ene reaction. A 20 kDa 8-arm PEG amine with a pentaerythritol core (JenKem) was functionalized with norbornene end groups via a HATU coupling. Detailed end-group functionalization procedures have been outlined previously by Ohnsorg *et al.*^19^. The ratio between thiols and norbornenes in the hydrogel was kept stoichiometric (1:1). SH-PPEGA9-100-SH (2.47 mM) was combined with PEG-NB (0.43 mM) in PBS with 1 mM CRGDS and 6.8mM LAP. The desired number of cells were added to the mixture prior to pipetting the solution into a mold affixed to a thiolated glass coverslip and irradiating with 365 nm light (3.5 or 5 mW/cm^2^) for 5 minutes. The mold was removed prior to transferring the hydrogel embedded with cells to culture media. The cells were then cultured within the hydrogels for up to 28 days changing media every other day (Schematic S2 and Figure S3).

#### Collagen Type-1 Hydrogels

All solutions used to formulate the collagen type-1 hydrogels were prepared on ice. The collagen stock solution was diluted to the desired concentration (3 mg/mL) using 10X PBS. The collagen solution was then neutralized using 1N NaOH, the solution was vortex, the cells were added and vigorously mixed via pipetting. Once mixed, the solution was pipetted into a trans-well insert and incubated at 37 °C for 30 minutes. Media was then added to the transwell and the cells were cultured for up to 28 days changing media every other day.

### Rheological Characterization

All rheological measurements were conducted on a TA Instruments DHR-3 Rheometer equipped with an 8 mm parallel plate (BB hydrogel) or a 20 mm parallel plate (Col1 hydrogel) geometry (either smooth or sandblasted). Experiments were either conducted on a quartz plate attached to a UV curing stage or a Peltier plate for temperature control. Each measurement was performed with at least n = 3 samples and the average was plotted along with the standard deviation.

#### In Situ Gelation

The bottlebrush polymer hydrogels were photopolymerized on a UV curing stage equipped with a quartz plate and using an 8 mm sandblasted upper geometry. An Omnicure 1000 light source (λ = 365 nm, *I* = 5 mW/cm^2^) was connected to the UV curing stage via a light guide. Oscillation fast sampling measurements were collected for 10 minutes at 1% strain and 1 rad/sec to measure the storage and loss modulus as a function of time. The UV light was turned on 30 seconds into the measurement and turned off after 5 minutes of light irradiation.

The collagen type-1 hydrogel was polymerized using a Peltier plate and 20 mm upper geometry. Immediately following neutralization of the collagen type-1, the solution was pipetted onto a Peltier plate set to 4 °C. The collagen gelation (storage and loss moduli) was measured over 30 minutes at 1% strain and 1 rad/sec. After loading, the temperature of the Peltier plate increased to 37 °C at a rate of 5 °C/minute and held at 37 °C for the remainder of the measurement.

#### Stress-Relaxation

Following *in situ* gelation on the rheometer, a stress-step experiment was initiated where 10% strain was applied to each sample over a 5 second induction period. The resulting oscillation stress was then measured as a function of time over 11 minutes for at least 3 individual hydrogels.

#### Strain-Sweep

Following *in situ* gelation, an oscillatory strain sweep, from 0.1 – 200%, was applied to each sample while measuring the storage modulus at a constant frequency of 1 rad/s. The calculated stress was then plotted as function of applied strain. The derivative of this curve yields the differential modulus (K’) which can be plotted as a function of the oscillation stress to visualize the critical stress of the hydrogel.

### Laser Scanning Confocal Imaging

After the desired number of days in culture, the cells within either BB or Col1 scaffolds, were prepared for live imaging experiments (vide infra) or fixed using 10% formalin for 30 minutes followed three 10-minute washes with DPBS. All confocal imaging experiments were performed using a Nikon AX R attached to a Ti2-E base with either a 20X (NA = 0.75) air or 20X long working distance (NA = 0.95) water immersion objective. Live imaging experiments were conducted using environmental control (37 °C, 5% CO_2_, 95% humidity).

### Fluorescence Recovery After Photobleaching (FRAP)

FRAP was performed on cells after 28 days of culture in either OM or GM to demonstrate cell network connectivity. If a gap junction inhibitor was used, 100 µM of CBX was added to culture approximately 24 hours before FRAP measurements. On day 28, cells were incubated with 1 µM Calcein AM in phenol free LG DMEM at 37 °C, 5% CO_2_, for 30 min. Samples were then washed with PBS for five minutes to remove excess Calcein AM. After which, samples were moved to a glass bottom well plate with 200 µL of phenol free LG DMEM with or without 100 µM CBX. Live cell FRAP experiments were performed on a Nikon AXR Lazer Scanning Confocal Microscope with a 20X air objective. Three to five isolated and connected cells were imaged in each gel (n = 2-3). An ROI was manually drawn around the selected cells, a z-stack taken 15 µm above and below the selected cell, the Calcein AM was bleached within the ROI (30 seconds at 100% laser power of 405 nm and 488 nm), and Calcein AM recovery was tracked for 10 minutes imaging every 30 seconds (30 um z-stack). Fluorescence recovery was quantified over time using FIJI. Grey values within ROI and of background were recorded using a sum slices z-projection over sampling duration. Background fluorescence was subtracted from ROI grey values. Normalized fluorescence recovery was plotted over time. Maximum recovery was recorded for each replicate.

### Connexin43 Immunostaining and Quantification

Before immunostaining, the fixed samples were permeabilized with 0.1% Triton X-100 for 1 hour at room temperature and blocked for 1 hour with 5% BSA at 4 °C. The primary antibody was then added in 5% BSA and incubated overnight at 4 °C [anti-Cx43 antibody, rabbit, (2.6 μg/mL or 1:200)]. The excess antibody was washed from the hydrogels using DPBS with 0.05% Tween 20 (PBST) for three, 10-minute washes. The secondary antibodies were then added in 5% BSA to the sample and incubated overnight at 4 °C [DAPI (1:500); Rhodamine Phalloidin (1:300); goat anti-rabbit Alexafluor 488 (1:500)]. The samples were then washed three more times for 10 minutes each wash with PBST before storing the samples at 4 °C until imaging.

Connexin 43 quantification was completed using FIJI. The f-actin signal was used to create a binary mask of the image to ensure that only connexin 43 puncta associated with the f-actin cytoskeleton were included in the particle analysis. A threshold was applied to the masked connexin 43 signal and particle analysis was used to measure the area and aspect ratio of the puncta. The particle analysis mask was then applied to an overlayed image of the nuclei and connexin 43 staining. From this image, the distance from each measured puncta to the nearest nuclei was measured. This analysis was conducted for at least three fields of view within each hydrogel (n = 3 hydrogels, each individual biological replicates).

### Calcium Transient Imaging and Quantification

After 28 days of culture, Fluo-8 AM (5 μM) and Pluronic F-127 (0.04 % w/v) were incubated with the cells in phenol-free LG-DMEM with 10% FBS for 30 minutes at 37 °C. After 30 minutes, the samples were then incubated for another 30 minutes at 25 °C before transferring the samples to a glass bottom 24 well plate with new Fluo-8 AM/Pluronic F-127, phenol-free LG-DMEM media for live imaging experiments on the laser scanning confocal microscope (Nikon AX R). At least 3 xyz positions were imaged per sample imaging z-stacks (50 μm; 5 μm z-step) every 20 seconds for 10 minutes.

The change in fluorescence intensity over time was quantified in FIJI from a sum slices z-projection. Drift correction (Fast4DReg) was first applied to ensure the cell of interest remained with the selected region of interest (ROI). Then the ROI manager was used to select at least 10 cells per image and an area of the background fluorescence intensity. The fluorescent intensity within the ROI was measured across all images in the acquisition using the Multi Measure function. The background fluorescent intensity was subtracted from the measured fluorescence intensity for all samples. The change in fluorescence intensity (ΔF = F – F_0_) was then normalized as a function of the baseline fluorescence intensity (F_0_) for each cell. A “flash” was quantified as an increase in greater than 200 fluorescence intensity units from one timepoint to the next. Average down sampling was applied to reduce the OM (n = 377) and GM (n = 260) to 100 measurements to best compare frequency of flashing cells between sample sets.

### Carbenoxolone Dose Optimization

hMSCs were seeded in a 96-well plate and cultured in either OM or GM. After 24 hours, varying concentrations of CBX (1-500 μM) were added to the respective media. After 24 hours of culture, a PrestoBlue assay was used to assess cell viability after treatment and the maximum dosage was chosen to be 100 μM (Figure S7) which was consistent with previous literature^24^.

### RNA isolation

RNA was isolated from the collagen and bottlebrush polymer hydrogels using the following methods. At least four hydrogels were pooled per RNA collection timepoint. The hydrogels were removed from trans-well plates or scrapped from glass coverslips into a 2mL microtube and flash frozen in LN_2_ with a 5 mm stainless steel bead. 1 mL of Trizol reagent was added to each sample and a bead mill tissue homogenizer was used to free the cells from the surrounding scaffold. The RNA layer was isolated from the Trizol reagent with chloroform. After phase separation via centrifugation, the top aqueous layer was removed and diluted with 70% ethanol. The RNA sample was then further cleaned and collected via a Qiagen RNeasy micro kit. Briefly, the sample was loaded onto an RNeasy MinElute spin column, rinsed with Buffer RW1, incubated with DNase, rinsed again with Buffer RW1, rinsed with Buffer RPE, rinsed with 80% ethanol, and eluted from the column using RNase-free water. RNA concentrations were immediately measured on a NanoDrop before storing at −80 °C until use.

### q-RT PCR

Reverse transcription was conducted using the SuperScript IV VILO Master Mix according to the supplier’s directions. The resulting cDNA was diluted 1:20 and stored at −20 °C until q-RT PCR was concluded. The cDNA, primers, and SYBR Green Supermix (Biorad) were combined and ran on the thermocycler according to the supplier’s directions.

## Acknowledgements

The imaging work was performed at the BioFrontiers Institute’s Advanced Light Microscopy Core (RRID: SCR_018302). The Nikon AXR Laser Scanning Confocal is supported by NIH Grant 1S10OD034320. The authors would also like to acknowledge McKenna Hanson and Brady Worrell at the University of Colorado Denver for use of their Tosoh EcoSEC with MALS system to characterize the bottlebrush polymer molecular weight. Additionally, the authors would like to acknowledge and thank Tamara Alliston for initial discussions that prompted first investigations into using the bottlebrush polymer scaffolds as a potential platform for 3D osteogenic differentiation. M.L.O acknowledges funding from NIH T32 AR080630. N.E.F. acknowledges funding from F32 AR084876. Finally all authors acknowledge funding from NIH R01 DE016523.

## Supporting Information

**Bottlebrush Polymer SEC and NMR and table of characterization.**

**Figure S1.**
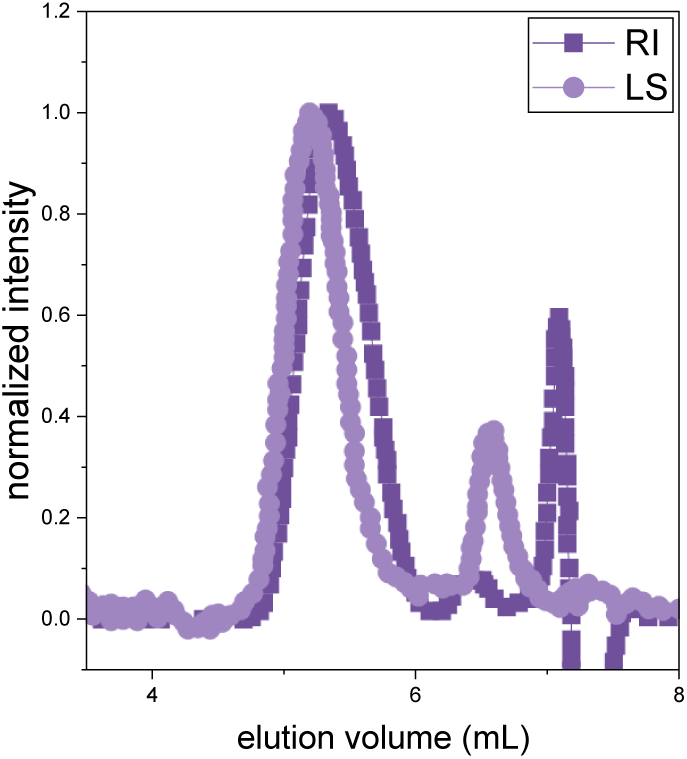
SEC MALS in DMF with 0.1% LiBr of the bottlebrush polymer crosslinker, C_12_H_25_-PPEGA9- **100**-C_12_H_25_.

**Table S1.** Molecular weight summary of the bottlebrush polymer crosslinker, C_12_H_25_-PPEGA9-100- C_12_H_25_.

|  | PPEGA9-**100** |
| --- | --- |
| $M_n$ | 22 kDa |
| $M_w$ | 48 kDa |
| $\bar{D}$ | 2.12 |
| dn/dc | 0.055 |

**Schematic S1.**
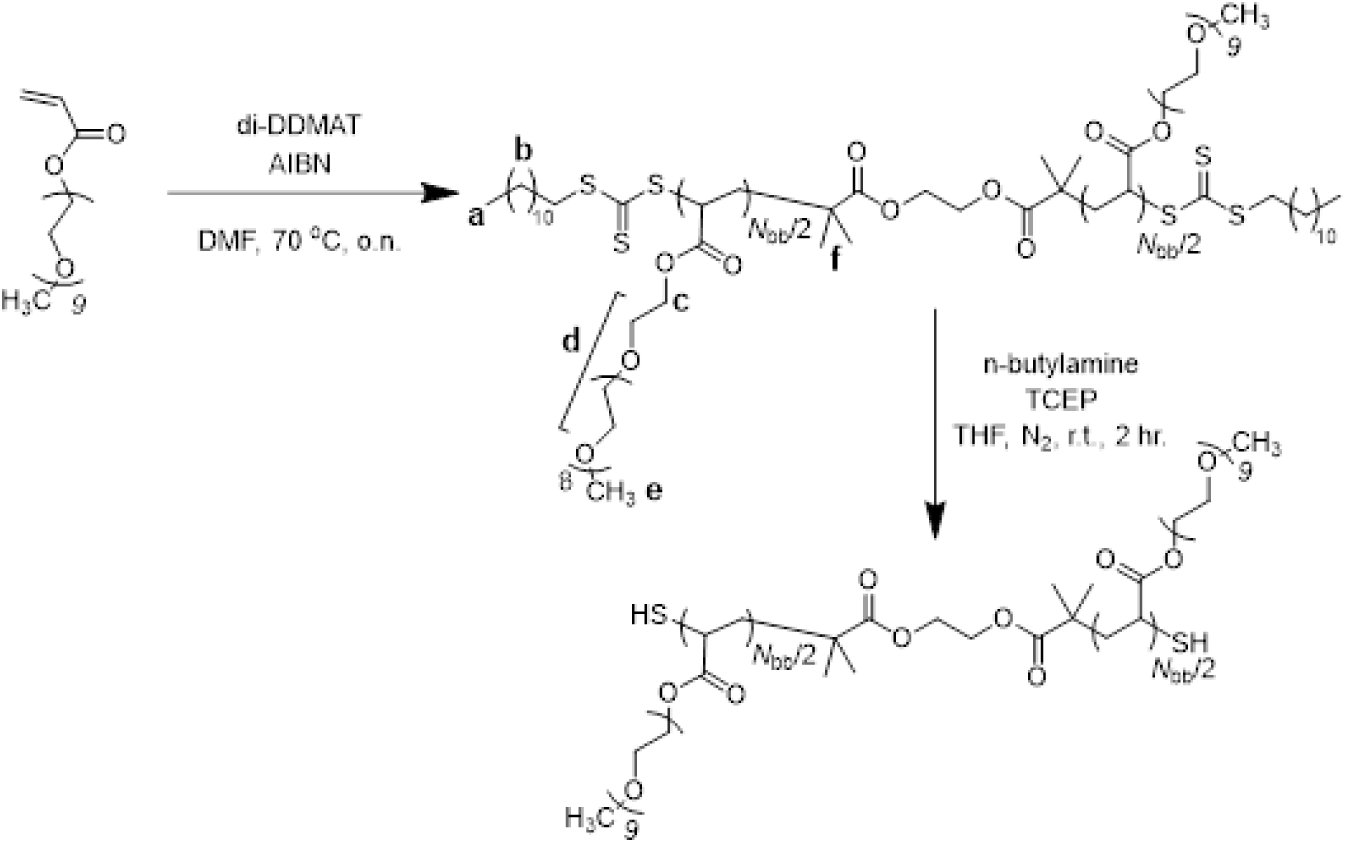
RAFT polymerization of the bottlebrush polymer and subsequent aminolysis to yield the bottlebrush dithiol crosslinker.

**Figure S2.**
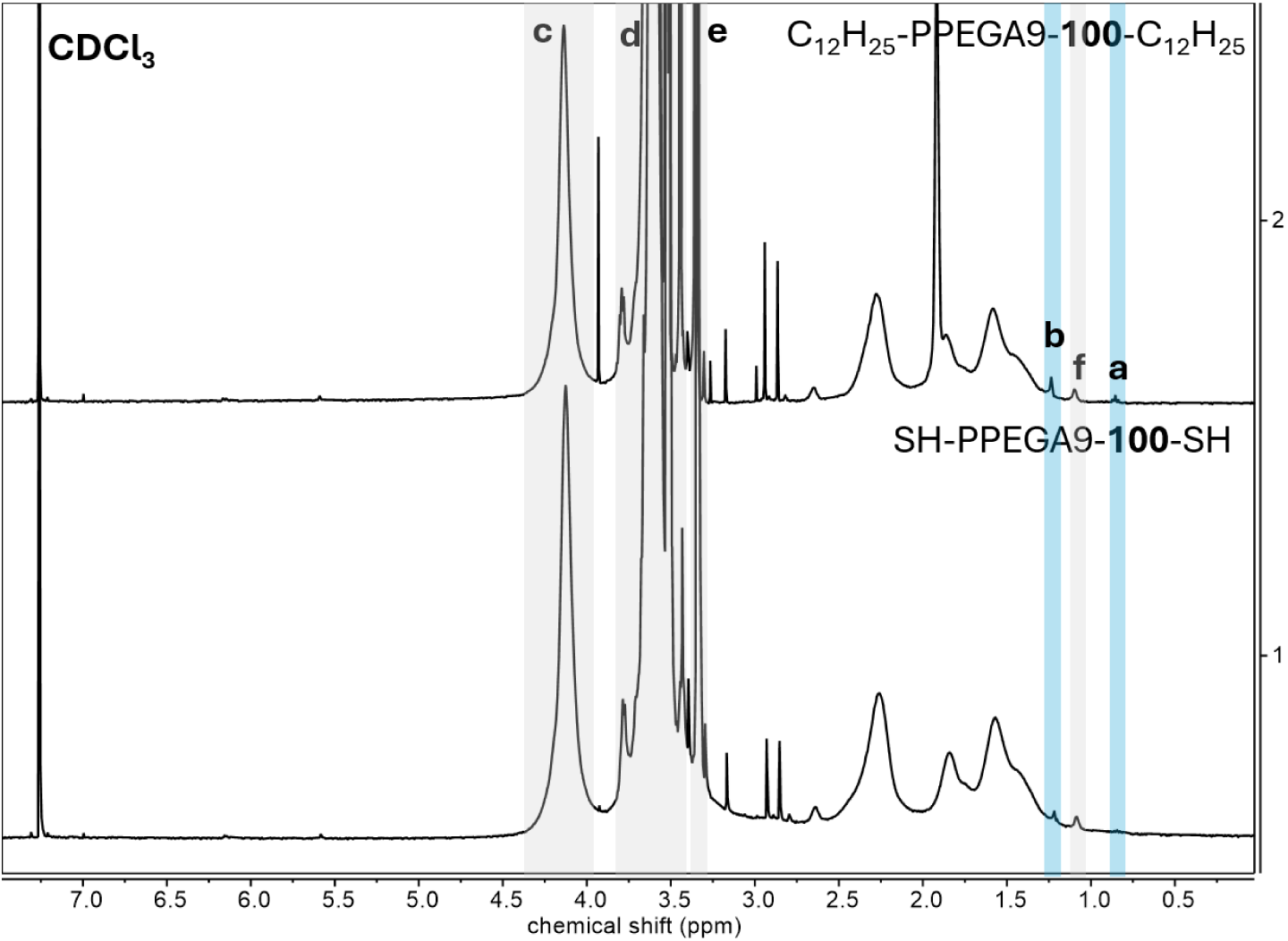
^1^H NMR of PPEGA9-98 before and after cleaving the trithiocarbonate end-groups.

**Schematic S2.**
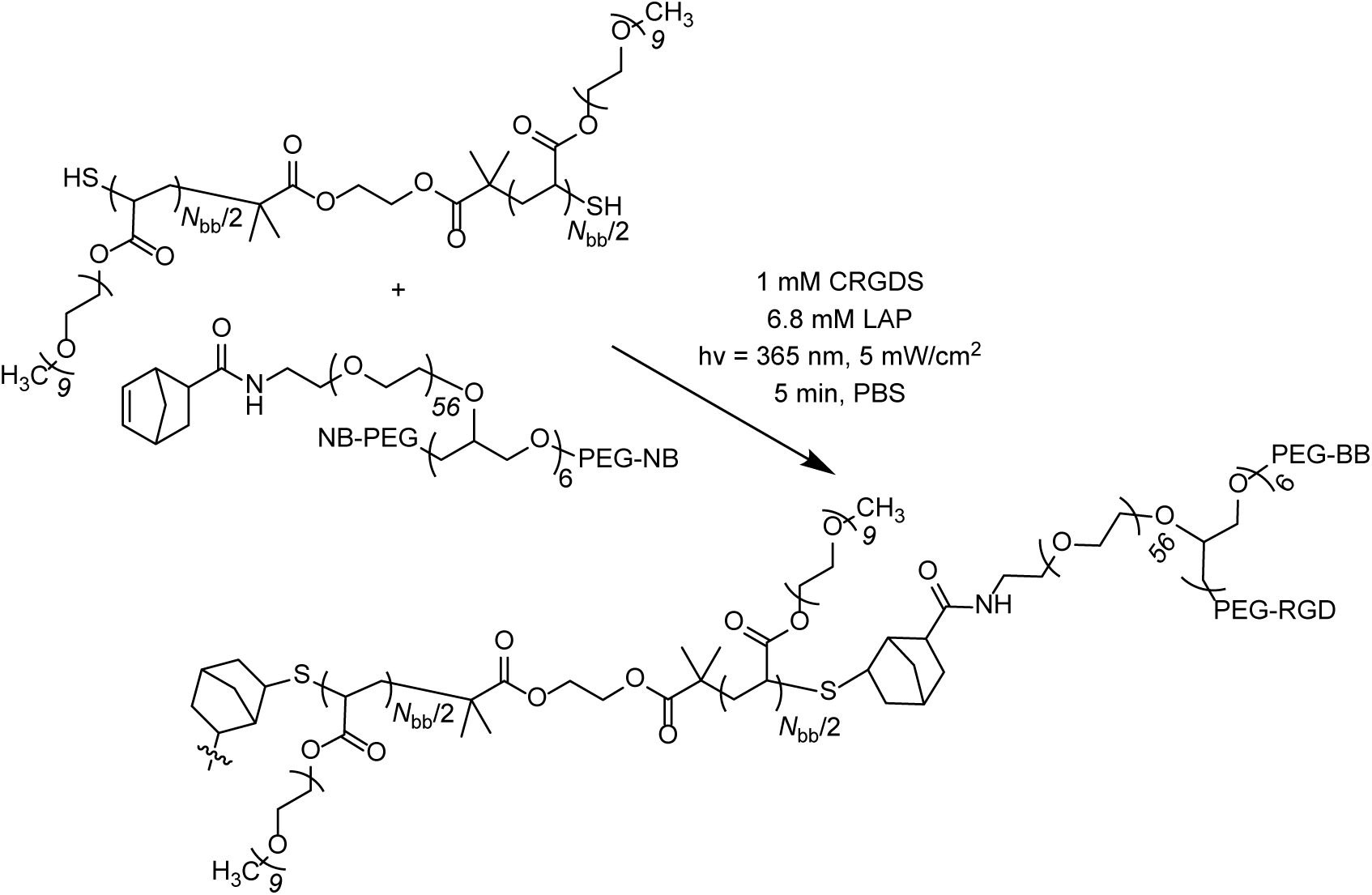
Thiol-ene photo-crosslinking of SH-PPEGA9-100-SH and an 8-arm PEG-norbornene (amide linked, pentaerythritol core) to form the bottlebrush polymer hydrogel network.

**Figure S3.**
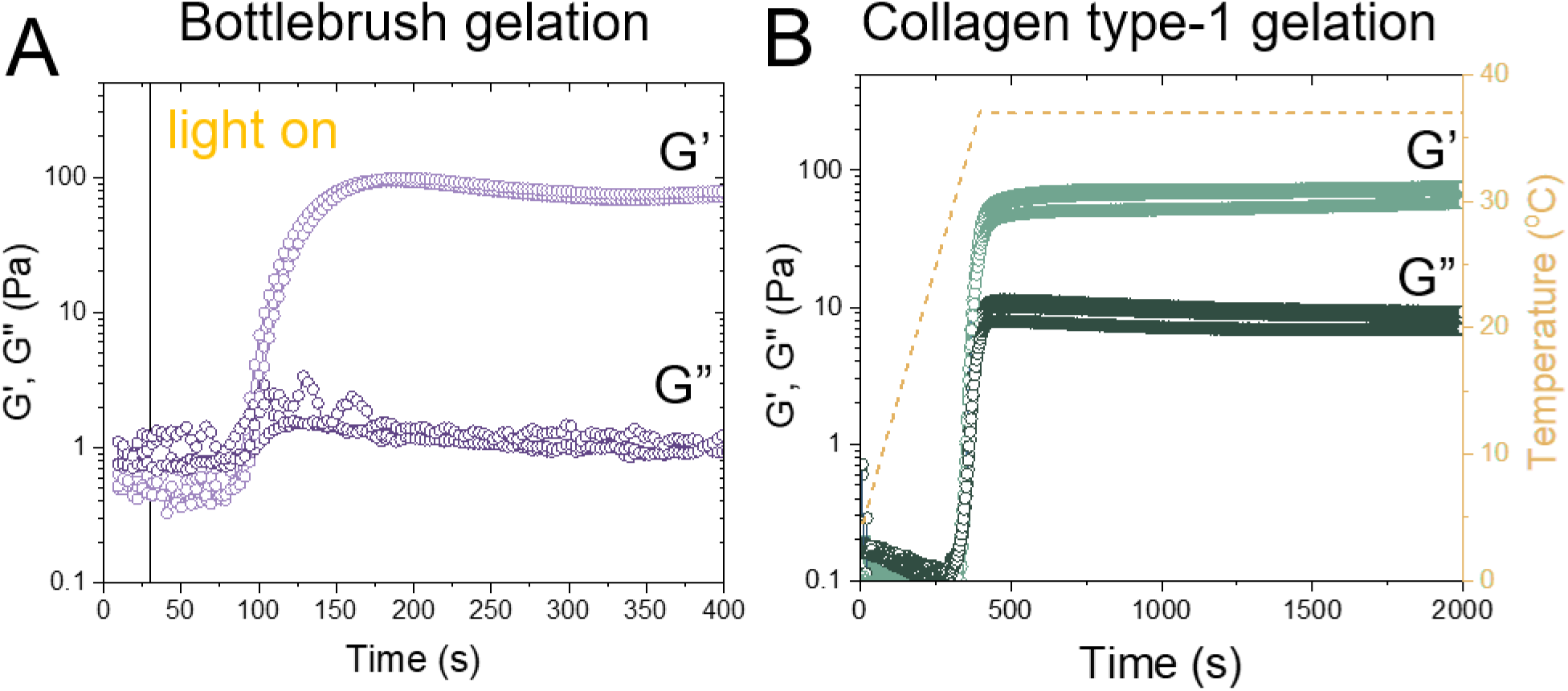
In situ rheology showing gelation of the (A) collagen gel under thermal gelation conditions and (B) the bottlebrush hydrogel undergoing photo-initiated crosslinking.

**Figure S4.**
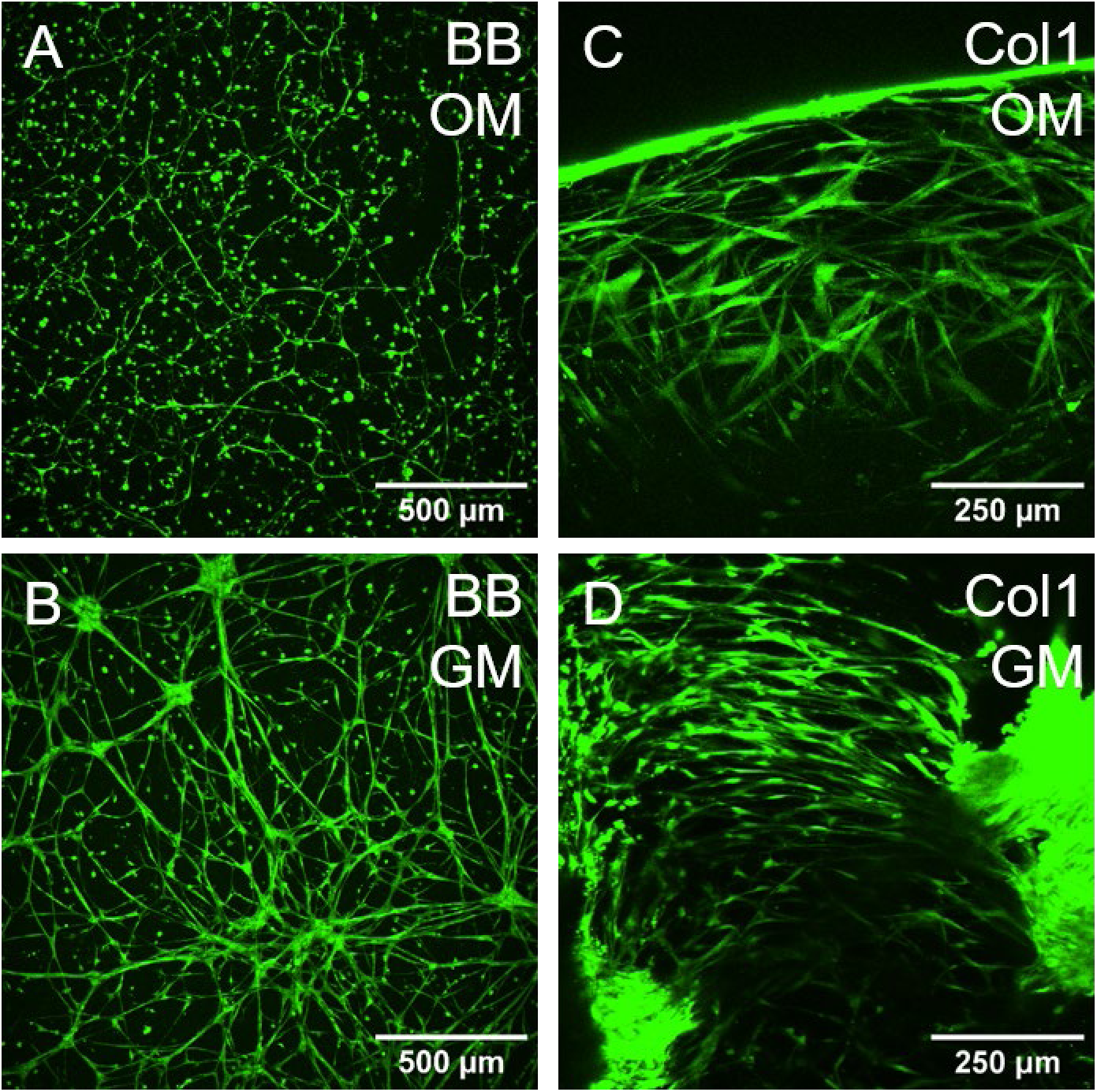
Live cell images (1 μM Calcein AM) showing majority of cells alive on day 28 for samples.

**Figure S5.**
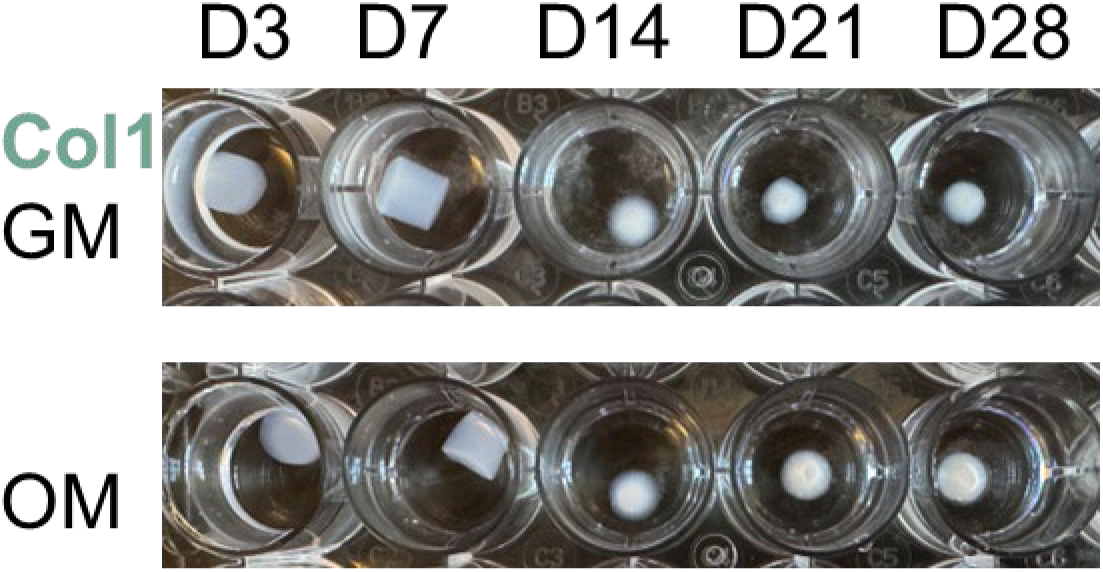
Cell-driven compaction of collagen type-1 hydrogels over the course of 28-days in culture. The gels were photographed within the wells of a 24-well plate after fixing and liberation from the trans-well insert.

**Figure S6.**
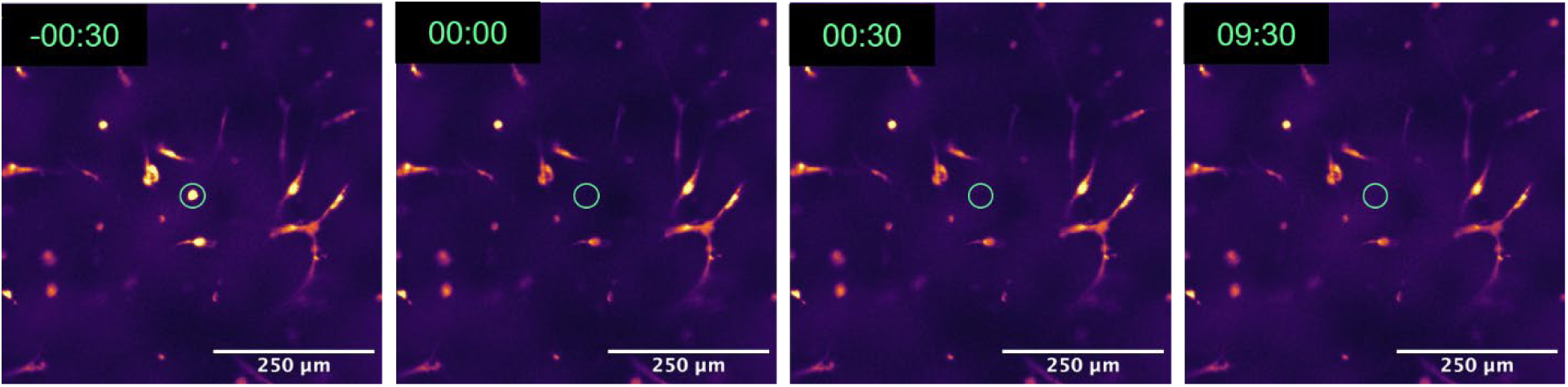
Image series showing a representative FRAP experiment on an isolated cell showing no fluorescence recovery over time within the ROI. −00:30 represents the fluorescence intensity pre-bleaching; 00:00 represents the total photobleaching of the Calcein AM within the ROI immediately following the photobleaching. The sample was then imaged every 30 seconds for 9 minutes and 30 seconds.

**Video S1.** Representative FRAP time-series of BB GM (AVI file)

**Video S2.** Representative FRAP time-series of BB OM (AVI file)

**Video S3.** Representative FRAP time-series of an isolated cell in BB OM (AVI file)

**Video S4.** Representative FRAP time-series of Col1 GM (AVI file)

**Video S5.** Representative FRAP time-series of Col1 OM (AVI file)

**Table S2.**
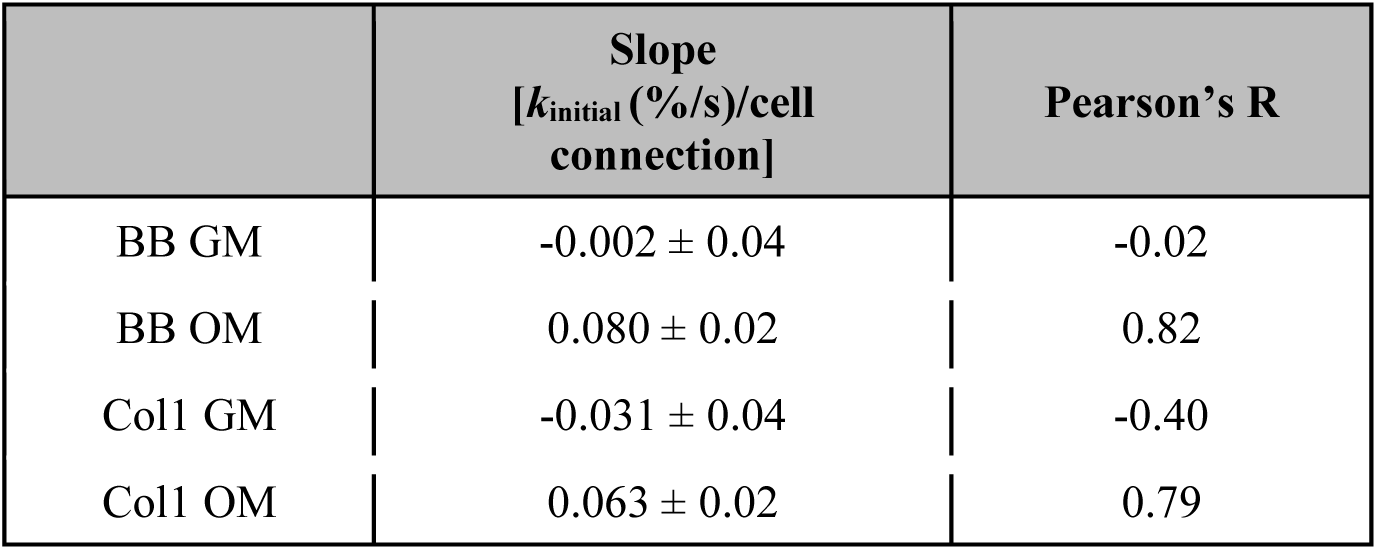
Summary of the initial rate of recovery plotted as a function of number of connected cells.

| | Slope<br>[$k_{\text{initial}}$ (%/s)/cell<br>connection] | Pearson's R |
| --- | --- | --- |
| BB GM | $-0.002 \pm 0.04$ | -0.02 |
| BB OM | $0.080 \pm 0.02$ | 0.82 |
| Col1 GM | $-0.031 \pm 0.04$ | -0.40 |
| Col1 OM | $0.063 \pm 0.02$ | 0.79 |

**Figure S7.**
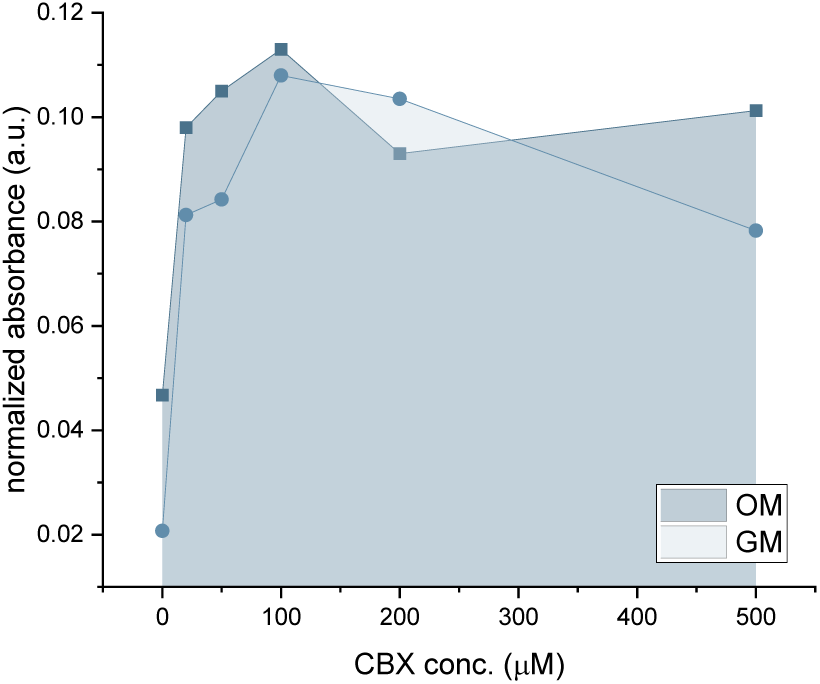
Carbenoxolone (CBX) dosing experiment to figure out maximum dosing concentration (100 μM) for GAP-FRAP studies.

**Video S6.** Representative GAP-FRAP time-series of BB GM (+) CBX (AVI file)

**Video S7.** Representative GAP-FRAP time-series of BB OM (+) CBX (AVI file)

**Video S8.** Representative GAP-FRAP time-series of Col1 GM (+) CBX (AVI file)

**Video S9.** Representative GAP-FRAP time-series of Col1 OM (+) CBX (AVI file)

**Video S10.** Representative Fluo-8 AM time-series of BB GM (AVI file)

**Video S11.** Representative Fluo-8 AM time-series of BB OM (AVI file)

**Video S12.** Representative Fluo-8 AM time-series of BB OM (dendrites) (AVI file)

## Notes

### Competing Interest Statement

The authors have declared no competing interest.

## References

1. Zaleske, MD, D.J. (2005). Embryology and Formation of Bone. In Netter’s Orthopaedics (Saunders), pp. 1–21.

2. Lin, X., Patil, S., Gao, Y.-G., and Qian, A. (2020). The Bone Extracellular Matrix in Bone Formation and Regeneration. Front. Pharmacol. 11. 10.3389/fphar.2020.00757.

3. McKee, T.J., Perlman, G., Morris, M., and Komarova, S.V. (2019). Extracellular matrix composition of connective tissues: a systematic review and meta-analysis. Sci. Rep. 9, 10542. 10.1038/s41598-019-46896-0.

4. Bonewald, L.F. (2011). The amazing osteocyte. J. Bone Miner. Res. 26, 229–238. 10.1002/jbmr.320.

5. Alliston, T. (2014). Biological Regulation of Bone Quality. Curr. Osteoporos. Rep. 12, 366–375. 10.1007/s11914-014-0213-4.

6. Sims, N.A., and Gooi, J.H. (2008). Bone remodeling: Multiple cellular interactions required for coupling of bone formation and resorption. Semin. Cell Dev. Biol. 19, 444–451. 10.1016/j.semcdb.2008.07.016.

7. Mazur, C.M., Woo, J.J., Yee, C.S., Fields, A.J., Acevedo, C., Bailey, K.N., Kaya, S., Fowler, T.W., Lotz, J.C., Dang, A., et al. (2019). Osteocyte dysfunction promotes osteoarthritis through MMP13-dependent suppression of subchondral bone homeostasis. Bone Res. 7, 34. 10.1038/s41413-019-0070-y.

8. Dole, N.S., Mazur, C.M., Acevedo, C., Lopez, J.P., Monteiro, D.A., Fowler, T.W., Gludovatz, B., Walsh, F., Regan, J.N., Messina, S., et al. (2017). Osteocyte-Intrinsic TGF-β Signaling Regulates Bone Quality through Perilacunar/Canalicular Remodeling. Cell Rep. 21, 2585–2596. 10.1016/j.celrep.2017.10.115.

9. Boukhechba, F., Balaguer, T., Michiels, J.-F., Ackermann, K., Quincey, D., Bouler, J.-M., Pyerin, W., Carle, G.F., and Rochet, N. (2009). Human Primary Osteocyte Differentiation in a 3D Culture System. J. Bone Miner. Res. 24, 1927–1935. 10.1359/jbmr.090517.

10. Zauchner, D., Müller, M.Z., Horrer, M., Bissig, L., Zhao, F., Fisch, P., Lee, S.S., Zenobi-Wong, M., Müller, R., and Qin, X.-H. (2024). Synthetic biodegradable microporous hydrogels for in vitro 3D culture of functional human bone cell networks. Nat. Commun. 15, 5027. 10.1038/s41467-024-49280-3.

11. Bernero, M., Zauchner, D., Müller, R., and Qin, X.-H. (2024). Interpenetrating network hydrogels for studying the role of matrix viscoelasticity in 3D osteocyte morphogenesis. Biomater. Sci. 12, 919–932. 10.1039/D3BM01781H.

12. Aziz, A.H., Wilmoth, R.L., Ferguson, V.L., and Bryant, S.J. (2020). IDG-SW3 Osteocyte Differentiation and Bone Extracellular Matrix Deposition Are Enhanced in a 3D Matrix Metalloproteinase-Sensitive Hydrogel. ACS Appl. Bio Mater. 3, 1666–1680. 10.1021/acsabm.9b01227.

13. Hoppock, G.A., Buettmann, E.G., Denisco, J.A., Goldscheitter, G.M., Condyles, S.N., Juhl, O.J., Friedman, M.A., Zhang, Y., and Donahue, H.J. (2023). Connexin 43 and cell culture substrate differentially regulate OCY454 osteocytic differentiation and signaling to primary bone cells. Am. J. Physiol.-Cell Physiol. 325, C907–C920. 10.1152/ajpcell.00220.2023.

14. Akiva, A., Melke, J., Ansari, S., Liv, N., Van Der Meijden, R., Van Erp, M., Zhao, F., Stout, M., Nijhuis, W.H., De Heus, C., et al. (2021). An Organoid for Woven Bone. Adv. Funct. Mater. 31, 2010524. 10.1002/adfm.202010524.

15. Caliari, S.R., and Harley, B.A.C. (2014). Structural and Biochemical Modification of a Collagen Scaffold to Selectively Enhance MSC Tenogenic, Chondrogenic, and Osteogenic Differentiation. Adv. Healthc. Mater. 3, 1086–1096. 10.1002/adhm.201300646.

16. Prince, E. (2024). Designing Biomimetic Strain-Stiffening into Synthetic Hydrogels. Biomacromolecules 25, 6283–6295. 10.1021/acs.biomac.4c00756.

17. Saraswathibhatla, A., Indana, D., and Chaudhuri, O. (2023). Cell–extracellular matrix mechanotransduction in 3D. Nat. Rev. Mol. Cell Biol. 24, 495–516. 10.1038/s41580-023-00583-1.

18. Xu, J., Jiang, Y., and Gao, L. (2023). Synthetic strain-stiffening hydrogels towards mechanical adaptability. J. Mater. Chem. B 11, 221–243. 10.1039/D2TB01743A.

19. Ohnsorg, M.L., Mash, K.M., Khang, A., Rao, V.V., Kirkpatrick, B.E., Bera, K., and Anseth, K.S. (2024). Nonlinear Elastic Bottlebrush Polymer Hydrogels Modulate Actomyosin Mediated Protrusion Formation in Mesenchymal Stromal Cells. Adv. Mater. 36, 2403198. 10.1002/adma.202403198.

20. Lund, A.W., Stegemann, J.P., and Plopper, G.E. (2009). Inhibition of ERK Promotes Collagen Gel Compaction and Fibrillogenesis to Amplify the Osteogenesis of Human Mesenchymal Stem Cells in Three-Dimensional Collagen I Culture. Stem Cells Dev. 18, 331–341. 10.1089/scd.2008.0075.

21. Plotkin, L.I., and Bellido, T. (2013). Beyond gap junctions: Connexin43 and bone cell signaling. Bone 52, 157–166. 10.1016/j.bone.2012.09.030.

22. Zappalà, A., Romano, I.R., D’Angeli, F., Musumeci, G., Lo Furno, D., Giuffrida, R., and Mannino, G. (2023). Functional Roles of Connexins and Gap Junctions in Osteo-Chondral Cellular Components. Int. J. Mol. Sci. 24, 4156. 10.3390/ijms24044156.

23. Liu, W., Cui, Y., Wei, J., Sun, J., Zheng, L., and Xie, J. (2020). Gap junction-mediated cell-to-cell communication in oral development and oral diseases: a concise review of research progress. Int. J. Oral Sci. 12, 17. 10.1038/s41368-020-0086-6.

24. Kuzma-Kuzniarska, M., Yapp, C., Pearson-Jones, T.W., Jones, A.K., and Hulley, P.A. (2014). Functional assessment of gap junctions in monolayer and three-dimensional cultures of human tendon cells using fluorescence recovery after photobleaching. J. Biomed. Opt. 19, 015001. 10.1117/1.JBO.19.1.015001.

25. Bonewald, L.F. (2013). Osteocyte Biology. In Osteoporosis (Elsevier), pp. 209–234. 10.1016/B978-0-12-415853-5.00010-8.

26. Kawano, S., Otsu, K., Kuruma, A., Shoji, S., Yanagida, E., Muto, Y., Yoshikawa, F., Hirayama, Y., Mikoshiba, K., and Furuichi, T. (2006). ATP autocrine/paracrine signaling induces calcium oscillations and NFAT activation in human mesenchymal stem cells. Cell Calcium 39, 313–324. 10.1016/j.ceca.2005.11.008.

27. Kawano, S., Shoji, S., Ichinose, S., Yamagata, K., Tagami, M., and Hiraoka, M. (2002). Characterization of Ca2+ signaling pathways in human mesenchymal stem cells. Cell Calcium 32, 165–174. 10.1016/S0143416002001240.

28. Qin, L., Liu, W., Cao, H., and Xiao, G. (2020). Molecular mechanosensors in osteocytes. Bone Res. 8, 23. 10.1038/s41413-020-0099-y.

29. Burra, S., Nicolella, D.P., Francis, W.L., Freitas, C.J., Mueschke, N.J., Poole, K., and Jiang, J.X. (2010). Dendritic processes of osteocytes are mechanotransducers that induce the opening of hemichannels. Proc. Natl. Acad. Sci. U. S. A. 107, 13648–13653. 10.1073/pnas.1009382107.

30. Storm, C., Pastore, J.J., MacKintosh, F.C., Lubensky, T.C., and Janmey, P.A. (2005). Nonlinear elasticity in biological gels. Nature 435, 191–194. 10.1038/nature03521.

31. Jaspers, M., Dennison, M., Mabesoone, M.F.J., MacKintosh, F.C., Rowan, A.E., and Kouwer, P.H.J. (2014). Ultra-responsive soft matter from strain-stiffening hydrogels. Nat. Commun. 5, 5808. 10.1038/ncomms6808.

32. Zhang, K., Barragan-Adjemian, C., Ye, L., Kotha, S., Dallas, M., Lu, Y., Zhao, S., Harris, M., Harris, S.E., Feng, J.Q., et al. (2006). E11/gp38 Selective Expression in Osteocytes: Regulation by Mechanical Strain and Role in Dendrite Elongation. Mol. Cell. Biol. 26, 4539–4552. 10.1128/MCB.02120-05.

33. Lu, X.L., Huo, B., Park, M., and Guo, X.E. (2012). Calcium Response in Osteocytic Networks under Steady and Oscillatory Fluid Flow. Bone 51, 466–473. 10.1016/j.bone.2012.05.021.

34. Wang, X., Huang, M., Tang, Y., Li, Y., Yang, Y., and Zhou, M. (2025). Hypotonic stimuli promote osteocyte dendrite formation by modulating actin dynamics via the TRPV4-CDC42 signaling pathway. Mater. Today Bio 34, 102120. 10.1016/j.mtbio.2025.102120.

